# Automated assignment of cell identity from single-cell multiplexed imaging and proteomic data

**DOI:** 10.1101/2021.02.17.431633

**Authors:** Michael J. Geuenich, Jinyu Hou, Sunyun Lee, Hartland W. Jackson, Kieran R. Campbell

**Author notes:** Authors contributed equally (Michael J. Geuenich, Jinyu Hou, Sunyun Lee).

## Abstract

The creation of scalable single-cell and highly-multiplexed imaging technologies that profile the protein expression and phosphorylation status of heterogeneous cellular populations has led to multiple insights into disease processes including cancer initiation and progression. A major analytical challenge in interpreting the resulting data is the assignment of cells to *a priori* known cell types in a robust and interpretable manner. Existing approaches typically solve this by clustering cells followed by manual annotation of individual clusters or by strategies that gate protein expression at predefined thresholds. However, these often require several subjective analysis choices such as selecting the number of clusters and do not automatically assign cell types in line with prior biological knowledge. They further lack the ability to explicitly assign cells to an unknown or uncharacterized type, which exist in most highly multiplexed imaging experiments due to the limited number of markers quantified. To address these issues we present Astir, a probabilistic model to assign cells to cell types by integrating prior knowledge of marker proteins. Astir uses deep recognition neural networks for fast Bayesian inference, allowing for cell type annotations at the million-cell scale and in the absence of previously annotated reference data across multiple experimental modalities and antibody panels. We demonstrate that Astir outperforms existing approaches in terms of accuracy and robustness by applying it to over 2.1 million single cells from several suspension and imaging mass cytometry and microscopy datasets in multiple tissue contexts. We further showcase that Astir can be used for the fast analysis of the spatial architecture of the tumour microenvironment, automatically quantifying the immune influx and spatial heterogeneity of patient samples. Astir is freely available as an open source Python package at https://www.github.com/camlab-bioml/astir.

## Introduction

Recent advances in high-throughput sequencing and imaging technologies have enabled high-dimensional profiling of gene expression, protein abundance, and phosphorylation status in both human tissues and model organisms. Much of the attention in computational methods development for these technologies has focused on single-cell RNA-sequencing (scRNA-seq), which has led to improvements in our understanding of organism development [1], drug resistance in cancer treatment [2], as well as the role of the tumour microenvironment in cancer initiation and progression [3, 4]. However, despite its ability to capture an unbiased representation of the whole-transcriptome at single-cell resolution, scRNA-seq cannot directly measure protein abundance or phosphorylation status and requires special modifications to operate on archival tissue [5], with spatial transcriptomics technologies such as HDST [6] suffering from similar drawbacks.

To address these issues, a range of technologies have been invented to measure the expression of multiple proteins while retaining the spatial location of origin in the tissue section, collectively termed “highly multiplexed imaging”. These include Imaging Mass Cytometry (IMC) [7] and Multiplexed Ion Beam Imaging (MIBI) [8], which stain tissues with antibodies conjugated with heavy metals followed by laser ablation and mass spectrometry, allowing for the unbiased expression quantification of around 40 features [9] while retaining the spatial location of origin of each cell. Additional related technologies such as MxIF [10] and CyCIF [11] have also been developed allowing for the multiplexed detection of proteins using microscopy and immunofluorescence techniques, thereby retaining spatial information while only requiring standard microscopy equipment.

A major analytical challenge when interpreting the large quantities of single-cell data generated by these technologies is to relate each high-dimensional cell phenotype to *a priori* known cell types, which is crucial for a number of applications such as linking tumour microenvironment composition to patient outcomes [12]. For single-cell protein and gene expression assays, several methods have been developed to assign cells to cell types that rely on clustering cells followed by the manual annotation of clusters [13–17] (Figure 1A). For lower-dimensional platforms such as flow cytometry, manual gating strategies are traditionally employed on expression values of marker genes. However, these methods require prior knowledge of marker gene expression thresholds, lack reproducibility [18], and cells that fall outside the thresholded values will not be assigned/be mis-assigned rather than being assigned probabilities that distribute across the most likely cell types. While these issues have recently been addressed in the field of scRNA-seq with the advent of computational tools such as CellAssign that use probabilistic modelling to identify cell types based on marker gene expression [12, 19], such tools have yet to be created for single-cell highly multiplexed imaging and proteomic technologies. An additional challenge for cell type assignment in these data modalities is the set of proteins measured is typically experiment-specific meaning supervised learning approaches to align cells to reference populations that have become popular in scRNA-seq analysis [13, 20] cannot be used.

**Figure 1:**
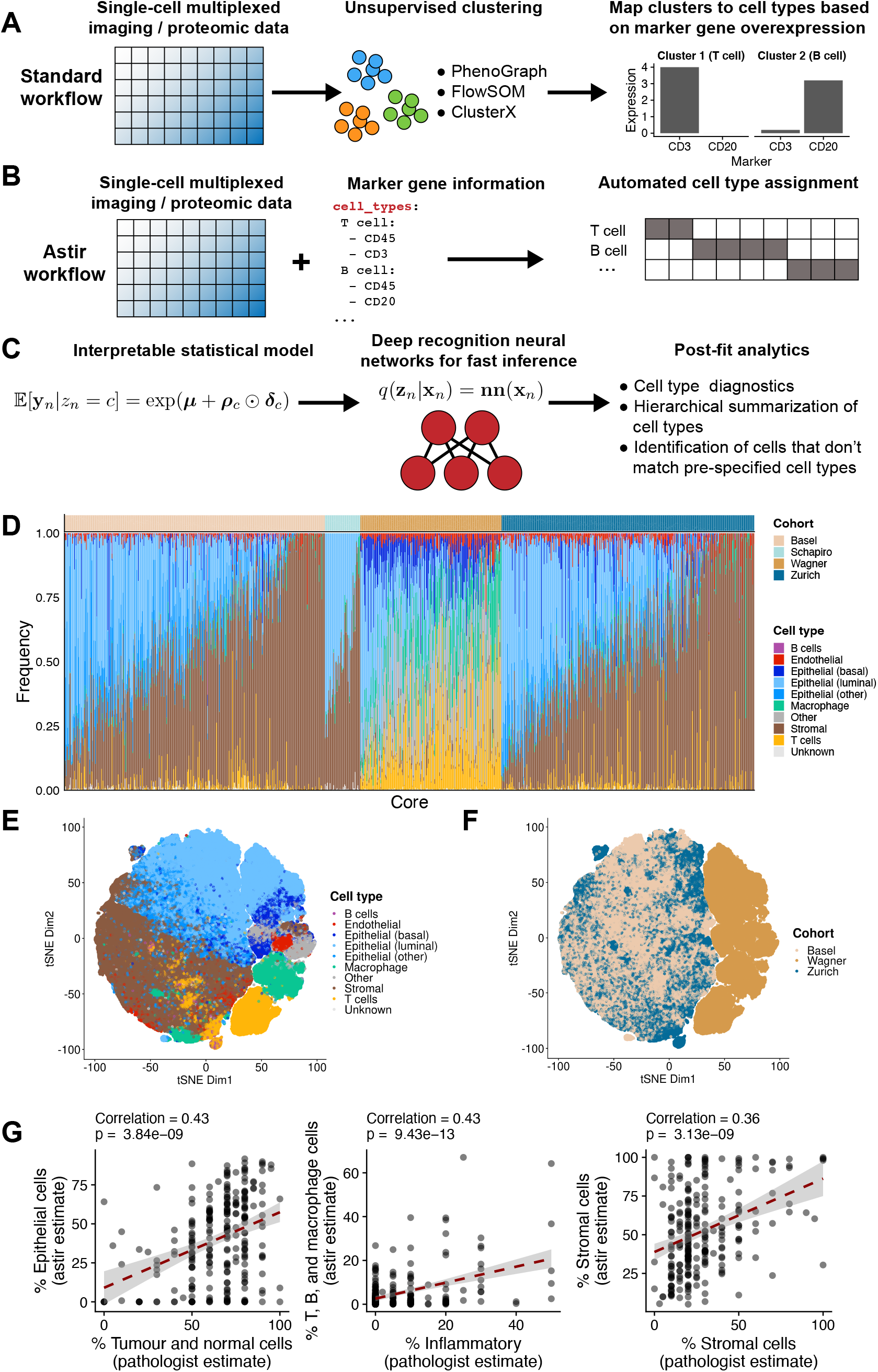
Astir automates cell type assignment for highly multiplexed imaging and single-cell proteomic data. **A** Existing cell type annotation methods cluster expression data and subsequently leverage marker over-expression to annotate clusters. **B** Astir combines input single-cell multiplexed imaging or proteomic expression data with a list of known marker proteins and probabilistically assigns each cell to a cell type. **C** Astir is based on an interpretable statistical model that assumes that marker proteins are more highly expressed in their respective cell types and uses deep recognition neural networks for fast Bayesian inference. post-fit diagnostics then allow for quality assessment of the assigned cell identities. **D** Astir identifies inter-patient tumour cellular composition heterogeneity across breast cancer samples from the Basel, Zurich, Schapiro, and Wagner cohorts. **E-F** tSNE plots for the Basel, Zurich and Wagner cohorts coloured by cell type and cohort demonstrates both cohort-specific variation and Astir’s ability to confidently assign cell types across cohorts. **G** A comparison of Astir cell type proportions to pathologist estimates (to the nearest 5%) based on immunohistochemistry for the Basel cohort demonstrates significant association across epithelial, inflammatory, and stromal compartments.

To address these challenges, we introduce Astir (ASsignmenT of sIngle-cell pRoteomics), a scalable probabilistic model to robustly assign cell types to data generated from a range of single-cell highly-multiplexed imaging and proteomic technologies. Expanding on our previous work [12], Astir takes both measured expression data and an *a priori* specified set of marker proteins for all expected cell types as input and employs a novel statistical machine learning model to assign cell type probabilities to each cell (Figure 1B). We apply Astir to three IMC datasets and one suspension mass cytometry dataset comprising a total of 552 patients and 930 cores from breast cancer tumours representing all clinical subtypes and grades of pathology, as well as normal tissue from the same patients. In addition, we include a CyCIF multiplexed microscopy dataset of formalin-fixed paraffin-embedded tissue comprising normal and cancer tissue from 40 different patients. We compare the Astir cell type assignment to multiple alternative workflows and show that Astir outperforms existing methods on average when assigning cells to the set of expected cell types present in the data and is uniquely able to assign cells to an *Other* type when cell type marker proteins are missing or unknown. We further demonstrate that Astir is more robust to variability in cell segmentation – a key upstream processing step in the analysis of highly multiplexed imaging data – and to imbalance in the underlying cell type proportions. We apply Astir to two IMC datasets of breast cancers and use the cell type assignments coupled with the spatial information to derive a novel interpretable spatial heterogeneity score that enables us to reproducibly track phenotypic changes with spatial tumour architecture across cohorts. Finally, we investigate the expression of marker proteins in *Unknown* and *Other* cells and show that this assignment is driven by the simultaneous expression of implausible marker proteins in part caused by non-specific staining of the individual cores.

## Results

### Automated assignment of cell type for single-cell multiplexed imaging and protein expression data

The Astir framework implements a statistical model that assumes a cell type represents a phenotypically static expression profile. Astir takes a cell-by-protein normalized expression matrix as input along with a yaml file specifying marker proteins for each expected cell type within the dataset, which is closely guided by the initial design of the protein measurement (antibody) panel. The Astir cell type model builds upon our previous work [12], assuming cell types express their marker proteins at relatively higher levels than other cell types with a modified likelihood function allowing for continuous intensity values as typically measured by such experimental technologies and a novel protein-protein covariance structure that requires marker protein expression to be correlated within each cell type (see *Methods*). It further allows for the existence of an *Other* cell type that does not express the specified marker proteins in a recognizable manner and may represent *de novo* cell types within the data. Further variation due to batch or patient specific effects may be incorporated in the model which supports the inclusion of arbitrary experimental design matrices.

To perform cell type assignment, Astir calculates the probability that each cell is of a given specified cell type under a Bayesian framework (see *Methods*). Notably, we employ deep neural networks [21] that map from the observed data to the cell assignment probabilities, allowing for fast mini-batch inference scaling to millions of cells that is both constant in memory and enables immediate prediction of cell types for new, unseen data (Figure 1C). To critique the resulting cell type assignments, Astir includes a set of post-fit diagnostics to ensure all cell types express their marker proteins at significantly higher levels than other cell types and flags all marker and cell type combinations for which this is violated. The Astir software is implemented in Pytorch [22], with a full set of documentation and tutorials online and is publicly installable from pypi^1^.

To demonstrate the benefits of Astir, we assigned cells to cell types across a number of highly-multiplexed imaging and suspension mass cytometry datasets. We collated (i) two cohorts of IMC data (“Basel” and “Zurich”) from a recent publication [23] of breast cancer tumour and normal samples comprising 307 patients and 735 cores with a total of 1.2 million cells, (ii) a suspension mass cytometry breast cancer dataset (“Wagner”) comprising 144 patients with one sample per patient and a total of 643,898 cells [24], (iii) a further IMC breast cancer dataset (“Schapiro”) with 51 patients and cores comprising a total of 45,585 cells [25], and (iv) a CyCIF multiplexed microscopy dataset from 13 different anatomical sites made up of cancer and normal tissue (“Lin”) comprised of 40 patients and a total of 179,558 cells [11]. Overall we assigned cell types to over 2.1 million cells using the marker proteins listed in Figure S1. Cells that coexpressed improbable marker combinations are automatically assigned as *Other* by Astir, while cells with a high probability of belonging to more than one cell type are assigned as *Unknown* (Figures S2 to S15).

Using Astir we assigned over 92% of cells to a cell type on average across all cohorts (Figure 1D), with the remainder assigned to *Unknown* and *Other* types. A certain fraction of cells being non-identifiable is expected given the limited set of proteins measured results in a fundamental inability to quantify all possible cell types that would require proteome- or transcriptome-wide measurement technologies. t-distributed stochastic neighbour embedding (t-SNE, [26]) reduced dimensionality representations of the cellular expression profiles based on overlapping proteins between the Basel, Zurich, and Wagner cohorts highlights a clear separation between the assigned cell types (Figure 1E) in the reduced dimensional space. It further shows separation between the Wagner and Basel/Zurich cohorts (Figure 1F) due to them originating not only from distinct experiments and antibody panels but from fundamentally different experimental technologies. This demonstrates a marked strength of Astir in its ability to automatically assign cells to common cell types across distinct experimental modalities.

Using the Astir cell type assignments we computed the proportion of each core containing a given cell type and compared to pathologist estimates of epithelial, stromal, and inflammatory compartments in cores from the Basel cohort. There was significant positive correlation in each case (Figure 1G), though with expected variation given the fundamental differences in measurement technologies. These largely arise due to pathologist estimates relying on cellular morphology with difficulty resolving small lymphocytes compared to the Astir assignments based on expression of marker proteins alone. We conclude that Astir can be used to robustly assign large numbers of cells to interpretable pre-defined cell types.

### Astir outperforms existing cluster-and-interpret methods and is robust to input marker mis-specification

We next compared Astir to existing workflows that assign cell types based on clustering followed by cluster interpretation via either inspection of marker proteins or gene set enrichment analysis. Using a set of commonly employed clustering methods for similar data types (PhenoGraph [15], FlowSOM [16] and ClusterX [17]), we clustered cells from all cohorts considered in an unsupervised manner across a range of plausible parameter values (*Methods*) and compared the identified clusters to the Astir assignments (Figures S16 to S45). Using the subsequently identified clusters, we sought to quantify how well each cluster corresponded to a desired cell type we wished to identify. Using gene set enrichment analysis [27], we matched each cluster to its most likely cell type and derived a score to quantify how well a given clustering can identify a cell type without further post-processing (*Methods*). Briefly, this awards each method a +1 if a given cell type within a dataset is resolved to exactly one cluster and – 1 if a given cell type is not identified or identified multiple times (i.e. corresponding to multiple clusters), with the sum of the individual scores representing the overall method score for a given dataset. We found that Astir outperformed all existing workflows across all cohorts (Figure 2A). This is largely driven by the fact that many methods will over or under cluster the given cells, even when the correct number of output cell types is specified (such as for FlowSOM k8 in the Basel cohort, see Figures S16 to S45 for representative examples).

**Figure 2:**
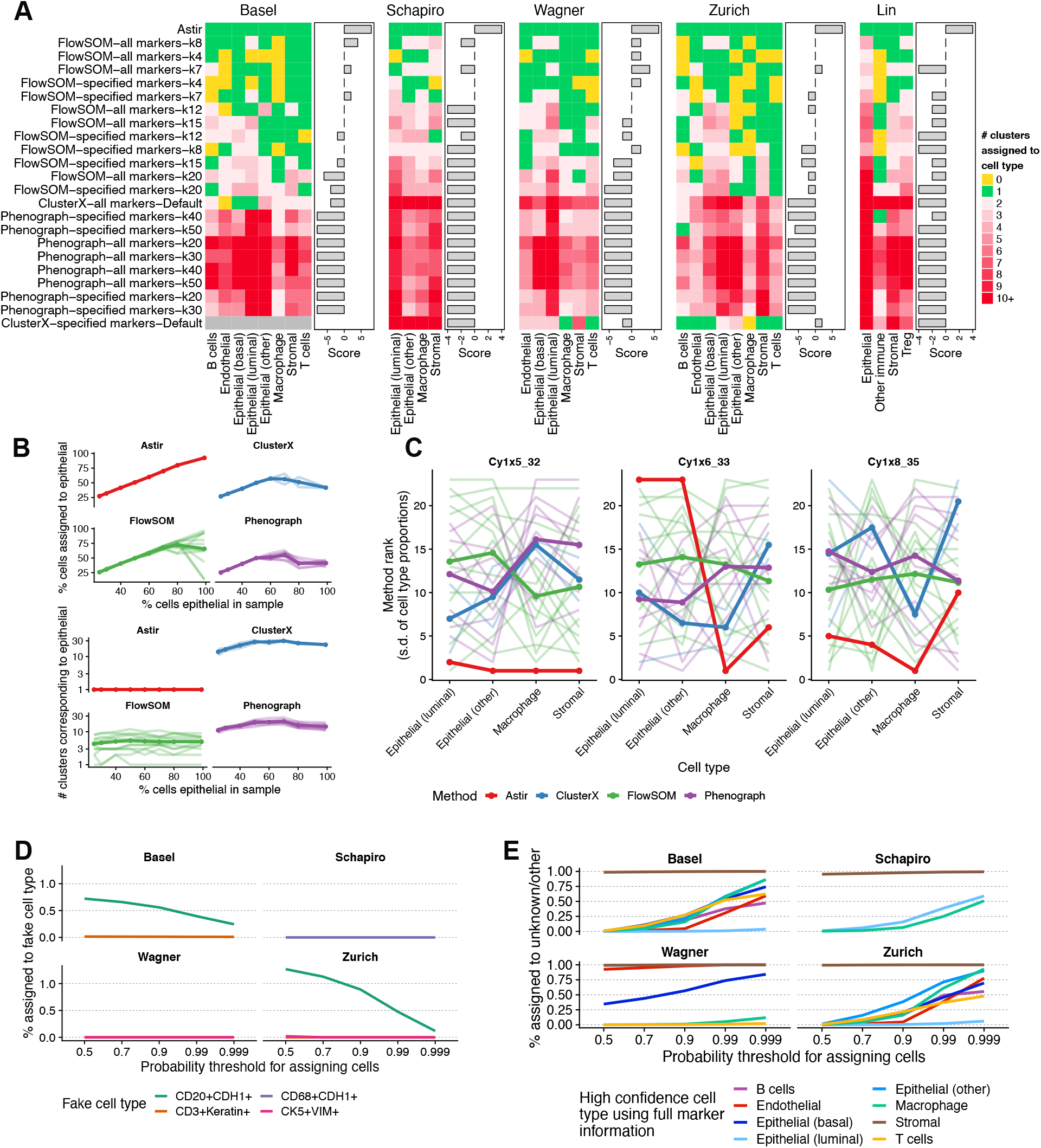
Comparison of Astir to existing workflows for cell type assignment and robustness to mis-specification of cell type marker proteins. **A** For each possible workflow (row), cell type (column) and dataset (individual heatmap), we quantified the number of clusters assigned to a given cell type. An ideal algorithm for cell type identification without further post-processing will assign a single cluster to each cell type (green), and not miss a cell type (yellow) or call multiple clusters per cell type (reds). Quantifying this as a “score” by adding a point if exactly one cluster is assigned per cell type and subtracting a point otherwise demonstrates Astir is the only method to assign cell types exactly. Only a single cluster was identified for ClusterX using markers specified in the Basel cohort resulting in the lack of an enrichment score. **B** Consistency between ground truth proportion of luminal epithelial cells in a simulated sample compared to inferred proportion (top) and number of clusters identified (bottom) across a range of methods. Each line represents a possible parameter configuration as per (**A**) with darker line representing the mean trend. **C** Astir has the highest overall consistency of cell type abundance estimates between different segmentations of iMC data across three regions of interest. The y-axis depicts the average rank of each method in terms of lowest standard deviation of cell type frequencies between segmentations. Each line represents possible parameter configurations as per (**B**). **D** percentage of cells that get assigned to a fictional cell type by Astir defined by combinations of unlikely marker combinations as a function of the probability threshold used to assign cells. **E** percentage of cells assigned as *Other* or *Unknown* upon the removal of markers for each cell type as a function of the probability threshold used to assign cells.

Next, we quantified the robustness of Astir and alternative methods to imbalance in cell type proportions in the input data. Taking 45,303 cells from the Basel cohort assigned to the luminal epithelial, stromal, macrophage, and T cell components with > 95% probability, we created new datasets by mixing the cells of different types with a pre-defined portion of luminal epithelial cells (25%, 30%, 40%….80%, 99%) with the other cells types being equally represented in the dataset. We then re-ran Astir and the alternative methods as above, and examined the impact of the underlying cell type proportion on both the percent of cells assigned to the luminal epithelial cell type as well as the number of clusters corresponding to luminal epithelial cells. We found Astir to be the only approach whose assignment of luminal epithelial cells matched the ground truth proportions (Figure 2B, top), with the existing approaches inferring incorrect proportions as the underlying composition became highly skewed towards an abundance of luminal epithelial cells. Similarly, we found Astir to be the only method where the number of luminal epithelial clusters inferred was independent of the underlying sample composition (Figure 2B, bottom). These results have important implications for the clinical utility of such methods given the importance of accurate cell type abundance identification to derive effective oncological biomarkers.

We next examined the robustness of each method to variability in cell segmentation of highly multiplexed imaging data. Cell segmentation is the process of taking the raw, pixel level data and computing the boundary of each cell via machine learning methods applied to membrane and nuclear markers [7] before summarizing to single-cell expression data. We leveraged existing data of three regions of interest (ROIs) segmented by three different users from a previous publication [25] to quantify the consistency of recovered cell type proportions from each method. We retrieved the segmentation mask for each user and ROI and summarized the cell specific expression as the mean expression of a given protein inside the cellular mask, and applied both Astir and the alternative methods to assign cell types. For each core, segmentation, and method, we computed the frequency of each cell type assigned and used the standard deviation in frequency across each segmentation as a metric, with a lower value implying more consistent assignment across segmentations.

Upon ranking each method we found Astir had the top overall ranked consistency between segmentations (Figure 2C), with each workflow based on clustering and post-hoc interpretation exhibiting large variations due with parameter configurations. An exception was the `Cy1×6_33` ROI where Astir ranked poorly in consistency of assignment across segmentations. Upon closer inspection, we found this to be caused by variation in assignment between `Epithelial (luminal)` and `Epithelial` (other) cell types (Figure S46), demonstrating variability in Astir’s cell type assignment in response to segmentation occurs between highly similar cell types. Overall, these results show Astir’s assignments are the most robust to upstream processing of highly multiplexed imaging data with important implications for a range of clinical applications.

We next benchmarked the robustness of Astir to mis-specification of the input cell type marker lists. We first created an input marker set containing biologically implausable (or “fictional”) cell types, including (i) co-expression of B-cell and epithelial markers (CD20_+_CDH1_+_), (ii) co-expression of T-cell and epithelial markers (CD3_+_Keratin_+_), (iii) co-expression of macrophage and epithelial markers (CD68_+_CDH1_+_, CD68_+_KRT6_+_), and (iv) co-expression of basal epithelial and fibroblast markers (KRT5_+_ViM_+_). We then re-assigned cell types using Astir including these fictional cell types in the input marker matrix for the four cohorts for which these markers were measured (Basel, Schapiro, Wagner, Zurich), at examined the proportion of cells erroneously assigned to these cell type as a function of the probability threshold for calling a given cell type (Figure 2D). We found that while the mis-classification rate is variable across both cohorts and cell types, Astir exhibits excellent specificity with no fictional cell type being assigned > 1% of cells when the probability threshold was moderately stringent (*p* ≥ 0.9).

Finally, we tested the robustness of Astir to the opposite scenario in which cell types present in the data are not specified in the input marker list. This is a fundamentally harder problem due to the correlated nature of expression data across cell types. As an example, consider the specification of a macrophage, typically defined by co-expression of CD45 and CD68. However, if we remove the canonical macrophage marker CD68 as input, cells instead simply appear as CD45-high, given CD45 is quantified for other cell types positive (such as T cells). Combined with non-specific staining of other markers, we therefore expect macrophage cells will be given some probability of being assigned as a T cell.

To quantitatively assess this, for every cohort we removed each cell type from the input set of cell types to which cells should be assigned and re-fit the Astir assignments. We then evaluated the proportion of cells newly assigned to *Unknown* or *Other* that were originally assigned to the removed cell type with high confidence (*p* > 0.95). The results show significant variability across both cohorts and cell types (Figure 2E). For example, for both the Wagner and Zurich cohorts, Astir demonstrates high sensitivity assigning almost 100% of cells originally assigned to Stromal as *Unknown /Other* when Stromal is removed from the input marker lists, presumably due to its phenotypic distinctiveness from other cell types present. In contrast, other cell types such as epithelial and T cells are erroneously assigned to alternative cell types, likely due to marker gene overlap with alternative epithelial and immune cell subsets.

We note that while Astir exhibits variability in this sense, the majority of existing methods do not have the ability to assign cells to *Unknown* and *Other* types. To quantitatively demonstrate this, we iteratively removed cell type markers starting with stromal cells, stromal and macrophages and finally, stromal, macrophage and endothelial cells prior to running the gene set variation analysis. This led to the misclassification of all stromal, macrophage and endothelial cells upon removal of their respective markers for all methods, parameters and cohort combinations (Figures S47 to S53). While it is theoretically possible to come up with a threshold for the GSVA score at which point a cluster is determined to be of an *Unknown* cell type, this is hard to implement in practice given that this threshold is not immediately obvious as the GSVA score does not represent a probability [27]. Thus, we conclude that Astir is uniquely capable of correctly assigning cells even when appropriate cell identity markers are unavailable, missing, or mis-specified.

### Interpretable stratification of patients based on cell type spatial heterogeneity signatures

We next sought to demonstrate that the retained spatial locations of cells coupled with the Astir cell type assignments could enable automated single-cell pathology and stratification of patients based on higher-order spatial features including immune cell influx or lack thereof. We calculated pairwise distances between all cell types in a given core and derived a core-specific spatial heterogeneity signature as the first principal component of the core-by-cell-type-interaction matrix (*Methods*). This creates a spatial heterogeneity score with higher values corresponding to smaller distances between different cell types (Figure 3A), enabling discrimination between mixed and compartmentalized spatial architectures, which have previously been associated with patient outcomes in triple negative breast cancer [28]. Indeed, inspection of cell types cores with the highest and lowest spatial heterogeneity scores identifies cases with extreme stromal-epithelial-immune mixing (case SP41_83_X14Y2, Basel cohort, Figure 3B left) and extreme compartmentalization between epithelial and T cells (SP43_114_X15Y1, Basel cohort, Figure 3B right).

**Figure 3:**
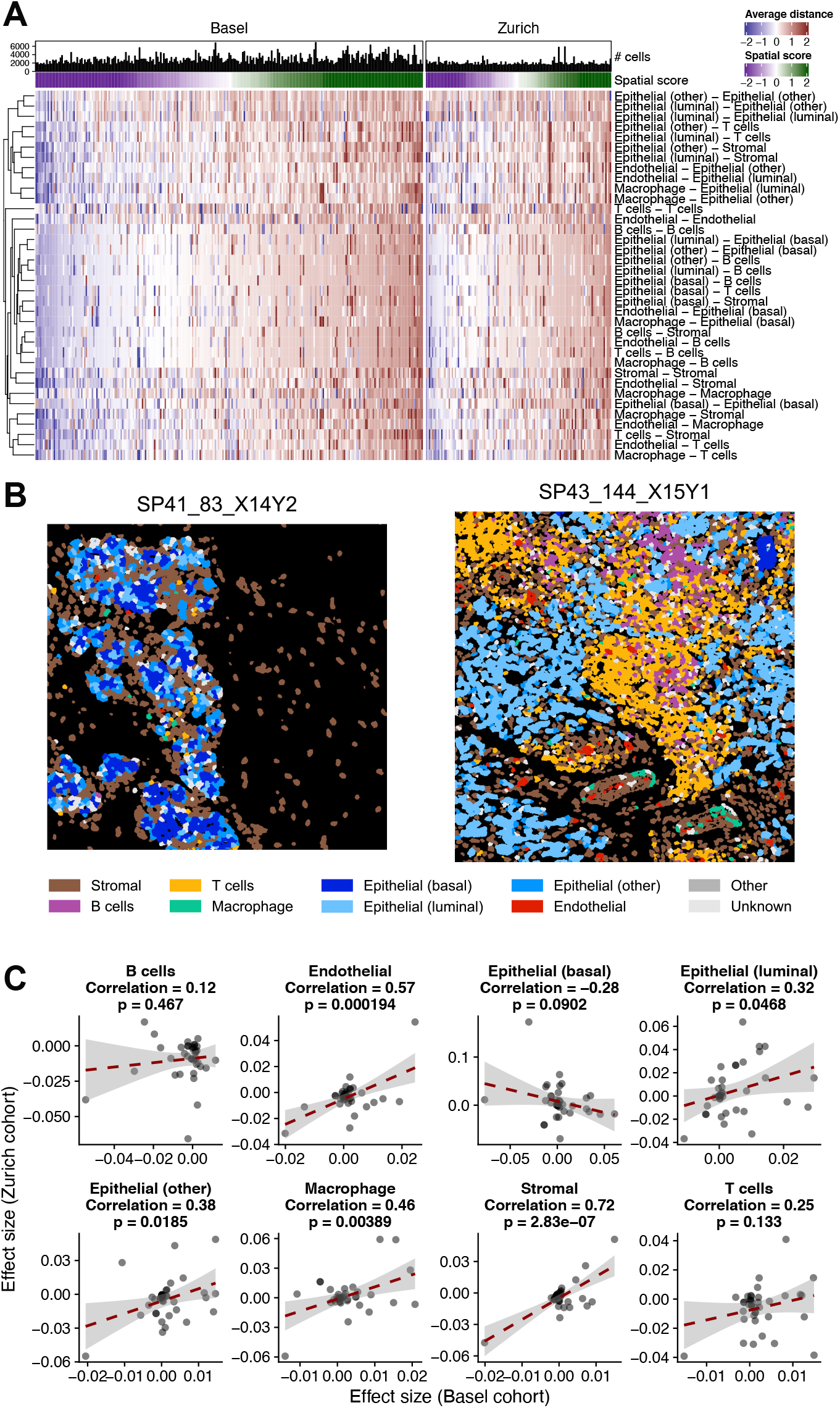
Automated quantification of immune influx and spatial heterogeneity from IMC identifies reproducible phenotypic perturbations across cohorts. **A** The average distance between each cell type pair per core tracks with the inferred spatial heterogeneity score. **B** Two representative cores with a low spatial heterogeneity score (left) indicating high levels of inter-cell type mixing, and a high spatial score (right) indicating a highly compartmentalized phenotype, coloured by Astir cell type assignment. **C** By regressing the per-core mean expression of each protein on the core’s spatial heterogeneity score in a cell type specific manner, we can compare the effect sizes between cohorts. This leads to a significantly correlated effect sizes between cohorts (Basel and Zurich), implying reproducible phenotypic changes associated with spatial tumour architecture.

Given the clinical importance of this spatial heterogeneity across a range of cancers, we examined whether the the impact of spatial tumour architecture on cellular phenotypes could be inferred in a manner reproducible between cohorts. To assess this, for each cell type we regressed the mean expression (per-core) of each protein on the spatial heterogeneity measure assigned to that core and compared the resulting effect sizes between each cohort (Basel and Zurich). The results were reproducible between cohorts, with 7/8 cell types displaying a positive correlation between cohorts in differential gene regulation with 5/8 significantly positively correlated at *p* < 0.05 (Figure 3C).

This analysis is only possible due to the systematic assignment of cell types by Astir that is reproducible between cohorts, enabling the comparison of a phenotypic profile of a given cell type between perturbative conditions such as tumour spatial architecture. This is in contrast to existing approaches whose clusters may conflate the notion of a cell type and phenotypic state, thus prohibiting the standardized measurement of distances between cell types without further post processing. Given the prognostic value of immune influx into the tumour mass [28], computational methods such as Astir that systematically assign cell types without further tuning will form the basis for standardized inference of complex tumour-immune architectures that can act as biomarkers in immuno-oncology.

### Identification of *Unknown* and *Other* cell types

Finally, we investigated the expression profiles of cells assigned as *Unknown* or *Other*. Given that the focus of highly multiplexed imaging experiments is often a heterogeneous population of cells and it is impossible to realistically design an antibody panel for all possible cell types, we expect any assignment cells to include those assigned to both *Unknown* and *Other*. However, we wished to ensure that the cell types classified as *Unknown / Other* do not over-express a particular combination of measured proteins, implying either we have mis-specified the input marker list or the Astir cell type assignments are incorrect.

Astir classified 0.7%, 0.8%, 25.4%, 0.4% and 3.4% of cells as *Unknown* or *Other* for the Basel, Schapiro, Wagner, Zurich and Lin cohorts respectively using the default setting that if a cell has > 50% probability of assignment it is assigned to a cell type. We conducted a differential expression analysis of all available measured proteins for each cohort, contrasting *Other* and *Unknown* cells against all assigned cells (Figure 4A-E). We found that across all cohorts, no proteins were expressed higher in *Unknown* and *Other* cells than in assigned cells with the exception of the stromal marker vimentin in the Wagner and Lin cohorts. Concerned that this may imply a mis-classification of stromal cells as *Unknown /Other*, we next contrasted the expression of all *Unknown / Other* cells against cells assigned as stromal, and found significantly decreased stromal marker expression and up-regulation of epithelial cell markers such as pan-Cytokeratin (Figure 4A-E).

**Figure 4:**
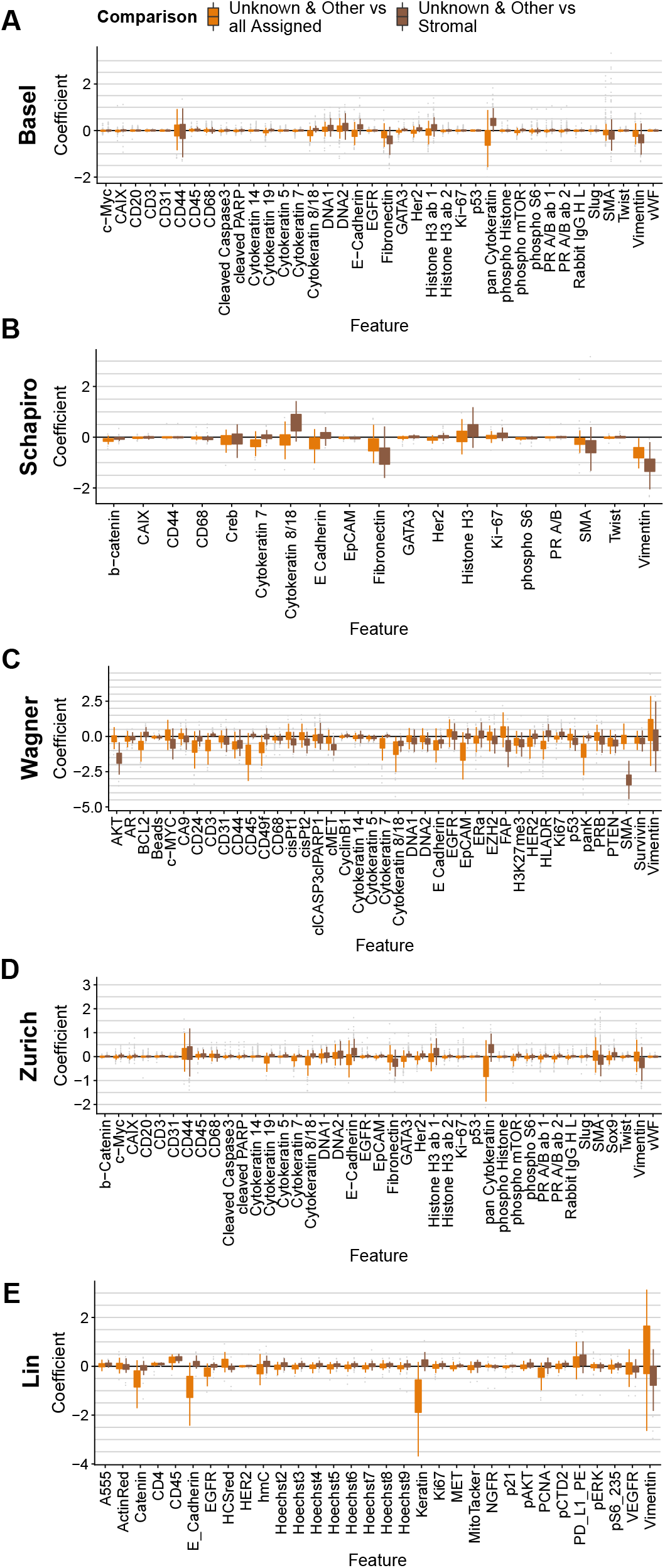
Analysis of *Unknown* and *Other* cells. Boxplots showing the distribution of the estimate of a linear model contrasting the expression of each protein in *Unknown* and *Other* cells compared to all assigned cells, and all *Unknown* and *Other* cells compared to stromal cells for the Basel (**A**), Schapiro (**B**), Wagner (**C**), Zurich (**D**), and Lin (**E**) cohorts.

Together, this analysis suggests those cells classified as *Unknown / Other* are not a distinct cell type identifiable from over-expression of a particular marker combination, but could instead represent an intermediate stromal-epithelial state, or is due to either mis-segmentation of neighbouring cells or non-specific antibody staining. To investigate whether the latter possibility was responsible for the assignment of *Unknown* and *Other* cells, we randomly selected a set of 20 cores from the Basel and Zurich cohort and coloured each image by the maximum assignment probability (Figure S54). Visual inspection indeed demonstrates general lower assignment probabilities at the tissue edge which could be related to reduced antibody staining quality and which we consequently conclude is one factor influencing the assignment of cells to *Unknown / Other* types by Astir.

## Discussion

Here we have introduced Astir, a novel probabilistic method for highly multiplexed imaging and single-cell proteomics designed to classify single cells into distinct cell types based on prior information of relevant marker proteins. While we have applied Astir to three IMC, one suspension CyTOF, and one CyCIF datasets, Astir may equally be applied to other expression assays including MIBI [8] and highly-multiplexed FISH. To our knowledge, Astir is the first dedicated tool for systematic and scalable assignment of cell types in these data modalities, where the use of classical referenced-based supervised learning approaches is not possible due to experiment-specific antibody panels.

As we have shown, an additional advantage of Astir is that the user can specify the probability threshold at which a cell is considered *Unknown* in type, thereby allowing for flexibility in the level of certainty required for the analysis at hand. Such an approach is not currently possible with any other method, which typically make a “hard” assignment of all given cells to a cell type regardless of accuracy. Similarly, while performing gene set enrichment analysis for the interpretation of clusters, it is theoretically possible to determine an enrichment score threshold at which a cluster is assigned to an *Unknown* type, identifying a relevant threshold that can be interpreted as a probability is challenging.

Although Astir is primarily designed for highly multiplexed imaging technologies, we do not currently make use of the spatial locations of cells for the purposes of cell type assignment. Future work may incorporate custom spatially-informed priors that have recently been incorporated into spatial clonal inference [29]. For example, in the context of breast cancers a cell adjacent to a large empty area interior to a duct (the lumen) is an epithelial cell. However, the spatial mapping of such anatomical structures is exceptionally challenging and we leave the incorporation of such anatomically-informed priors as future work.

As the volume of single-cell multiplexed imaging and proteomic data continues to grow, further phenotypic states will become apparent as both the number of cells and markers profiled increases. Consequently, we anticipate Astir becoming an additional and highly-used method in the interpretation of such data modalities for the discovery of novel cellular mechanisms, clinical biomarker studies, and personalized medicine.

## Methods

### Astir cell type model

The Astir cell type model requires two inputs: a protein expression matrix and a list of marker genes for the cell types of interest. The expression matrix is an *N* by *G* matrix of protein expression **Y** for *N* single cells and *G* proteins. These data could be generated by single-cell assays that measuring protein abundance, such as mass cytometry and imaging mass cytometry (post segmentation).

The list of marker genes for all cell types of interest is required as a *yaml* file but is re-coded into a binary matrix ***ρ*** where *ρ_gc_* = 1 if protein *g* is a marker for cell type *c* and 0 otherwise. Next, the conditional probability that a cell is of a certain type is specified based on the assumption that cell type marker proteins are more highly expressed in their respective cell types than other cells. Thus the expected expression of protein *g* in cell *n* given that that cell *z_n_* is of cell type *c* can be formulated as

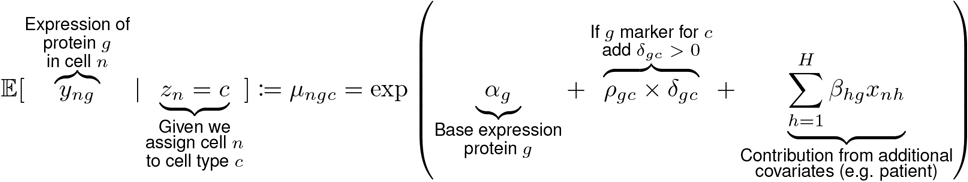

where *α_g_* is the base mean of protein *g* across all cell types and *δ_gc_* > 0 is the overexpression of protein *g* in cell type *c*. Therefore, when protein *g* is a marker for cell type *c*, the conditional expectation is modelled as being exp(*ρ_gc_* × *δ_gc_*) = exp(*δ_gc_*) times higher than in a cell type for which *g* is not a marker protein (i.e. ρ_gc_ = 0). Note that the overexpression parameter *δ* is cell type specific, allowing certain cell types to overexpress a given marker more than others (e.g. multiple types of macrophage overexpress *CD68* to different degrees). Finally, additional cell-specific covariates (such as cell cycle stage or patient of origin) may be incorporated through *H* covariates via the design matrix **X**.

Rather than assume the expression of marker genes are conditionally independent in a given cell type as in previous work [12], we introduce a novel co-variance structure that requires the expression of proteins to be correlated within a given cell type. Let **Σ**^(c)^ be the *G* × *G* matrix of pairwise covariances between proteins in cell type *c* which we specify as

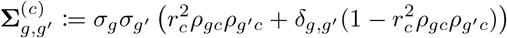

where *δ_g,g′_* = 1 if *g* = *g′*, and 0 otherwise.

This ensures the covariance has three desirable properties:

1. If *g* and *g′*, are both markers for cell type *c* and *g* ≠ *g′*, then the product *ρ_gc_ρ_g′c_* = 1 and the covariance between *g* and *g′*, is 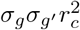, where *r* is a cell-type specific gene-gene correlation. Note we constrain *r* ∈ [0,1] since if two genes are a marker for a cell type they should not be anti-correlated.
2. If either *g* or *g′*, aren’t markers for *c* then and *g ≠ g′*, then *ρ_gc_ρ_g′c_* = 0 and the covariance is 0.
3. If *g* = *g′*, then the covariance becomes the variance 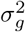

If we additionally define ***μ**_nc_* = [*μ_n1c_*,…, *μ_nGc_*] then we define the likelihood for a given cell *n* expression vector ***y**_n_* as

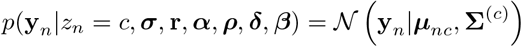

where 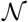 is the multivariate normal density. Then assuming the conditional likelihoods for each cell are independent we can write the likelihood for the entire expression matrix as

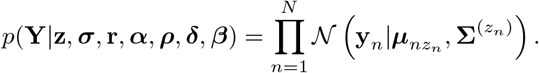

We append an additional column to *ρ* filled with constant 0, to signify the existence of an *Other* cell type that does not overexpress any marker and thus acts to capture all cells that do not fit a pre-defined cell type. Finally, we specify a prior for cell type assignment 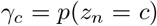 on the simplex such that 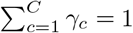 with a hierarchical prior specification *γ_1_*,…, *γ~* ∼ Dirichlet(*C*,…,*C*).

### Astir cell type inference

The overall object of inference is the posterior distribution of cell type assignments *p*(**z**|**Y**). Since this is impossible to calculate analytically, we resort to variational inference introducing variational distributions to minimize

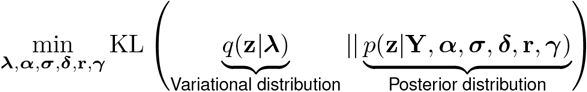

where KL is the Kullback-Leibler divergence. Rather than explicitly instantiate variational distributions *q*(*z_n_* = *c*) = *ψ_nc_* for each parameter, we use recent results from amortized inference [21] to propose a variational distribution of the form

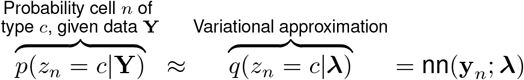

where nn is a neural network that maps the input data **y**_n_ for cell *n* to a simplex of probabilities over cell types, where *λ* are the (variational) parameters of the recognition neural network.

### Astir cell type implementation

Astir is implemented as a Python package using Pytorch for inference. Given the non-convexity of the minimization problem, the model is trained using five random initializations for a small number of epochs, following which the instantiation with the lowest ELBO is selected for continued training.

The cell type model parameters are initialized as follows:

- ***ρ*** is a matrix of size *G* × (*C* +1). Each entry *ρ_gc_* = 1 if protein *g* is a marker for cell type *c*, and *ρ_gc_ =* 0 otherwise.
- ***α*** is a vector of size *G*. To initialize ***α*** the following procedure is used: (i) a copy of the input expression matrix is created standardized to have empirical mean 0 and standard deviation 1, (ii) for each specified cell type *c* a set number *N_L_* of “putative cells” of type *c* is inferred as the *N_L_* cells with the highest standardized expression of the markers of cell type *c* with *N_L_* = 10 by default, (iii) for each protein *g*, the set of all cell types for which *g* is not a marker is collated, and the expression of *g* in the “putative cells” belonging to these cell types is averaged, with the log value of this average forming the initial value of *α_g_*.
- ***σ*** is a vector of size *G*. Element at *σ_g_* is initialized to be the standard deviation of expression value of protein *p* across all cells.
- ***δ*** is a matrix of size *G* × (*C* + 1). Each element at *δ_gc_* is the overexpression of protein *g* in cell type *c*. Its initial log value is sampled from normal distribution with mean 0 and variance 0.1
- ***r*** is a matrix of size *G* × (*C* + 1). It is a cell-type specific gene-gene correlation which is constrained within [0,1] and is initialized to be 0.5.
- The set of all parameters used in the recognition network including weights and bias are randomly initialized by PyTorch’s nn module. By default, the recognition network has two hidden layers of size 20 with a leaky ReLU activation [30].

### Diagnostics

The Astir package comes with a set of functions to perform post-fit diagnostics on the outputs of the model. This diagnostic ensures that all cell types express their marker proteins at significantly higher levels than other cell types, and flags all marker-cell types combinations for which this is violated. To identify cell types that violate this a t-test is run on the expression levels of a marker *g* between cells for which *g* is a marker and cells for which *g* is not a marker, and an error logged of the marker is not expressed significantly higher (at *α* = 0.05 significance) in the cell type for which it is a marker.

### IMC data processing & Astir cell type assignment

For the Basel and Zurich IMC cohorts, processed and spillover-corrected data were already available and normalized via an arcsinh transformation [31, 32] with a co-factor of 5 and winsorized to [0%, 99.9%]. For the Schapiro IMC cohort, the raw image files and masks were used to summarize to single-cell expression, where the expression of each protein in each cell was computed as the mean expression of that protein across all pixels belonging to that cell (code at https://github.com/camlab-bioml/schapiro-2017). The resulting data was normalized as per Basel and Zurich cohorts. Expression data from the Wagner cohort was similarly normalized with an arcsinh transformation and cofactor of 5. For the Lin cohort an arcsihn transformation with a cofactor of 100 was employed, with expression values winsorized to [1%, 99%]. The markers used by Astir to assign cell types for all cohorts are listed in Figure S1.

### Benchmarking Against Existing cell-type identification approaches

We compared the performance of Astir to that of FlowSOM [16], ClusterX [17] and Phenograph [15] using cytofkit [17], all of which are existing methods commonly used for the identification of cell types for flow and mass cytometry. For FlowSoM, cluster sizes were chosen to match the expected number of cell types (e.g. 8 for the Basel cohort) and varied such that potential cell states or Other cells could also be captured. For FlowSOM we used cluster sizes k = 4, 7, 8, 12, 15, and 20, for Phenograph we used nearest neighbour values 20, 30, 40 and 50, and for ClusterX we used default values.

Upon having assigned each cell to a cluster we used a gene set variation analysis [27] using the same cell type markers listed in Figure S1 to assign each cluster to a cell type. We did this by calculating the average enrichment for each cluster and cell type. For instance, for cluster one we calculated the average enrichment for T cells by computing the mean gsva enrichment across all cells in cluster one. We then repeated this for all target cell types and assigned the cell type that had the highest average enrichment to cluster one. We repeated this process for all clusters.

We compared the performance of the methods tested using an interpretability and accuracy score. Given that the true clusters are Unknown and that the goal of any cell type assignment is to generate a single interpretable assignment for each cell type, we created a score to measure the number of clusters per cell type. This score penalizes a method for assigning multiple cell types per cell type or assigning no clusters to a cell type by −1, and rewards a method for assigning exactly one cluster to each cell type with a score of +1.

### Spatial signature analysis

To identify novel spatial signatures enabled by Astir’s cell type assignment, we computed the mean distance from every cell type to every other cell type on the Basel and Zurich cohorts [23] by computing the pairwise distance from the centre of each cell for all possible cell type pairings followed by taking the mean. To enable distance comparisons across cell type pairs, the distances were standardized (mean 0 variance 1) in a cell-type-pair specific manner. Only cores with at least 1,000 cells present were retained to prevent low numbers of cells skewing distance calculations. Any missing distance values due to missing cell types from cores were imputed to the median of observed values for given cell-type pairs.

### Identification of *Unknown* cell types

To confirm the status of cells assigned as *Other* and *Unknown*, we conducted a differential expression analysis for the entire protein panel available for each cohort. We did this by fitting a linear model using the lm function in r to the expression of each protein and regressed on whether a cell was assigned as *Other /Unknown* or assigned as one of the specified cell types. We repeated this, splitting the data into *Other /Unknown* and Stromal cells and regressed on these categories as described above.

### Software packages used

With the exception of the single-cell spatial image plots which were created in Python using matplotlib [33] and seaborn [34] all plots were made using custom R scripts. tSNE plots were created with FasttSNE [35] and visualized using ggplot [36] and the wesanderson package for the cohort colour palette. All heatmaps were created using the ComplexHeatmap R package [37], alluvial plots were made with ggalluvial [38] and correlations were calculated with the Hmisc package. All scripts to recreate the analysis are available online at https://www.github.com/camlab-bioml/astir-manuscript.

## Data availability

The data for the Basel and Zurich cohorts used in this study are available through Zenodo at https://zenodo.org/record/3518284. The data for the Schapiro cohort are available at http://www.bodenmillerlab.org/research-2/histoCAT/. Masks corresponding to the different segmentatations are available upon request from the authors. The data for the Wagner cohort are available through Mendeley Data at https://data.mendeley.com/datasets/gb83sywsjc/1. The data for the lin-cycif cohort are available through the HMS LINCS publication page https://lincs.hms.harvard.edu/lin-elife-2018/.

## Author contributions

K.R.C. conceived the study and machine learning models. M.J.G. conceived the analysis plan, performed data analysis and created figures. J.H. and S.L. coded astir python package and performed additional analysis. M.J.G., H.W.J., S.L, J.H, and K.R.C. wrote the manuscript.

## Acknowledgements

We acknowledge the support of the Natural Sciences and Engineering Research Council of Canada (NSERC). This research was undertaken, in part, thanks to funding from the Canada Research Chairs Program.

## Supplemental figures

## Footnotes

1 https://pypi.org/project/astir/

## Notes

### Competing Interest Statement

The authors have declared no competing interest.

