## Supplementary figures for "Automated assignment of cell identity from single-cell multiplexed imaging and proteomic data"

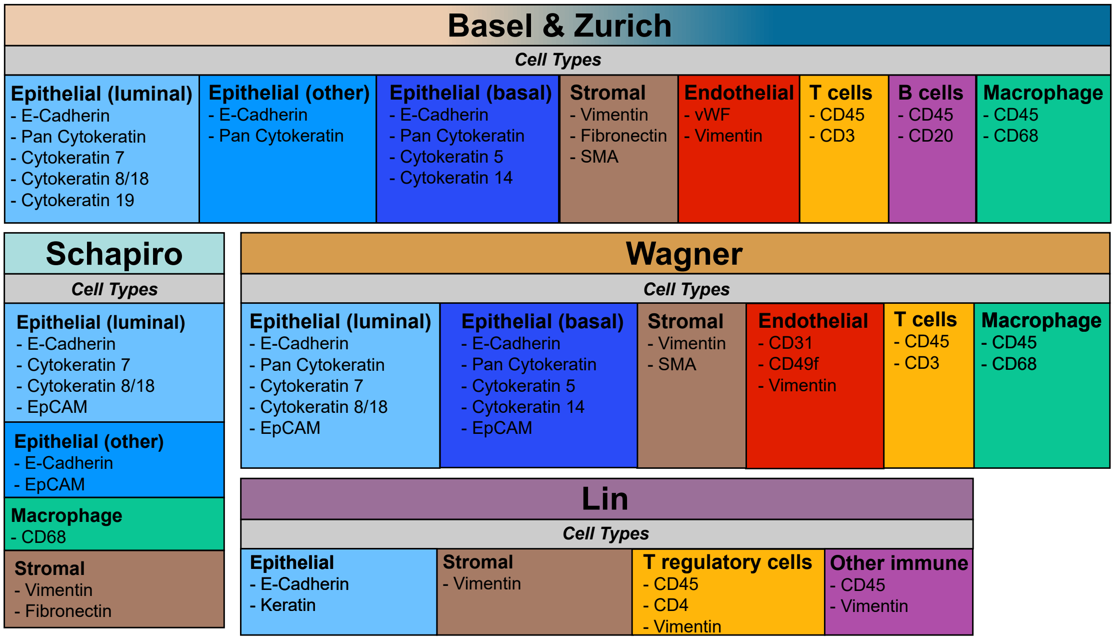


**Figure S1:** Cell type marker proteins.

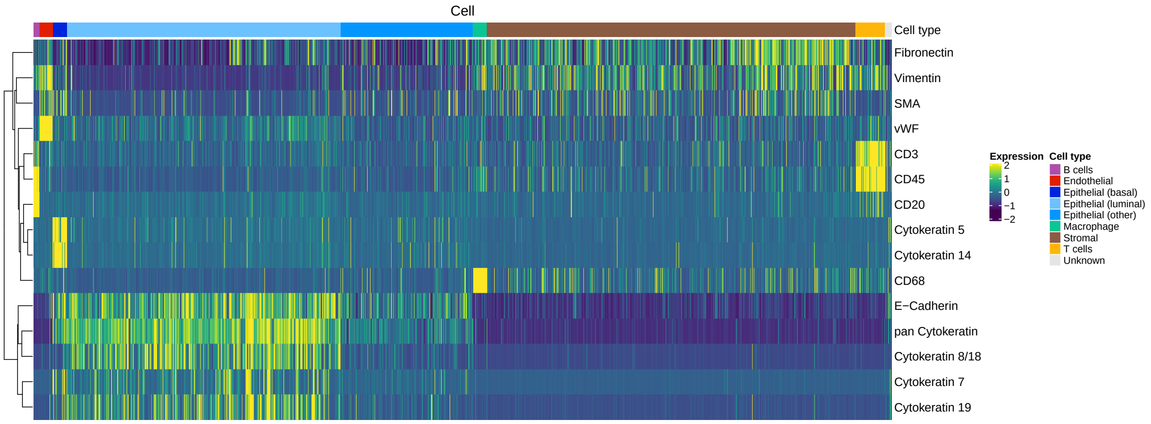


**Figure S2:** Expression heatmap for all marker proteins and the associated Astir cell types for Basel.

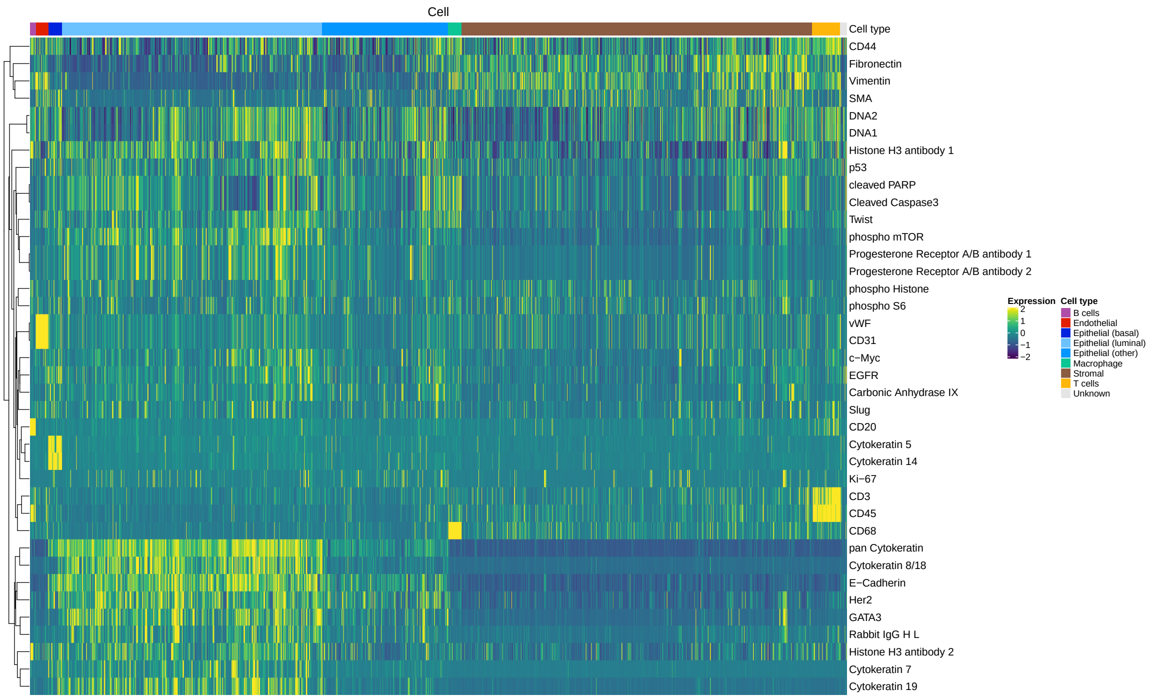


**Figure S3:** Expression heatmap for all proteins and the associated Astir cell types for Basel.

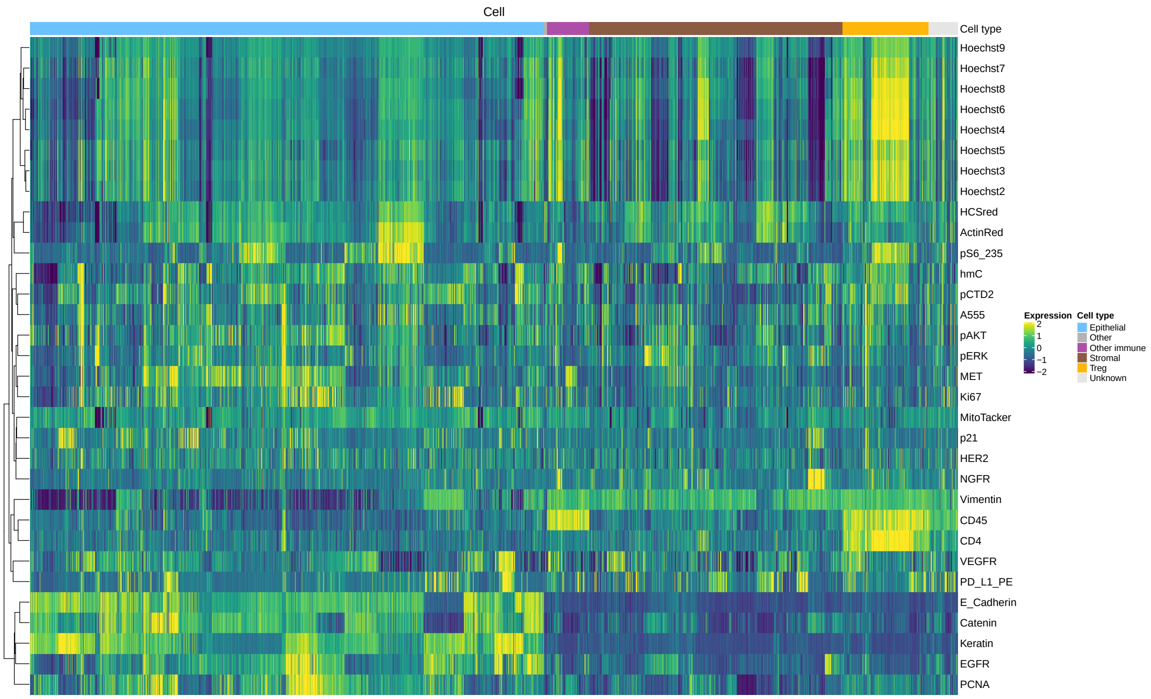


**Figure S4:** Expression heatmap for all proteins and the associated Astir cell types for Lin.

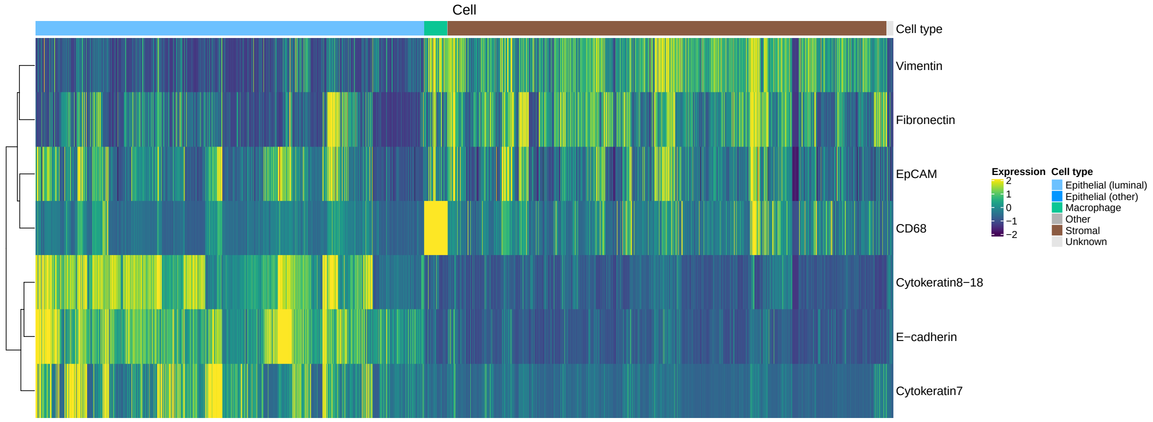


**Figure S5:** Expression heatmap for all marker proteins and the associated Astir cell types for Schapiro.

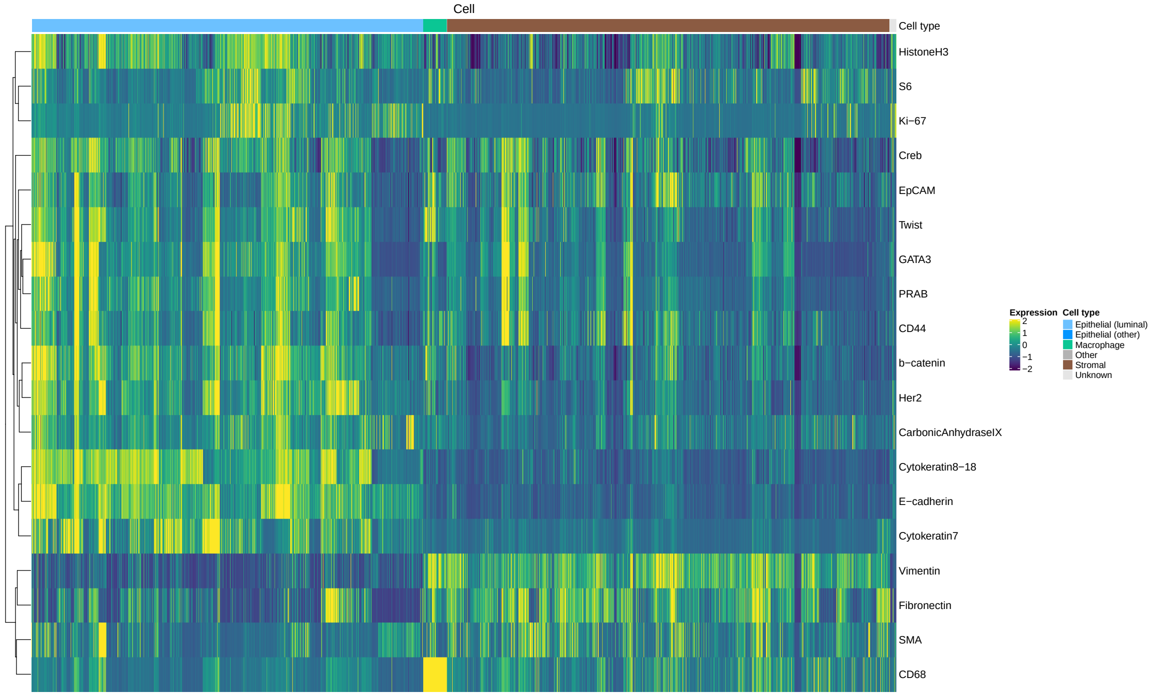


**Figure S6:** Expression heatmap for all proteins and the associated Astir cell types for Schapiro.

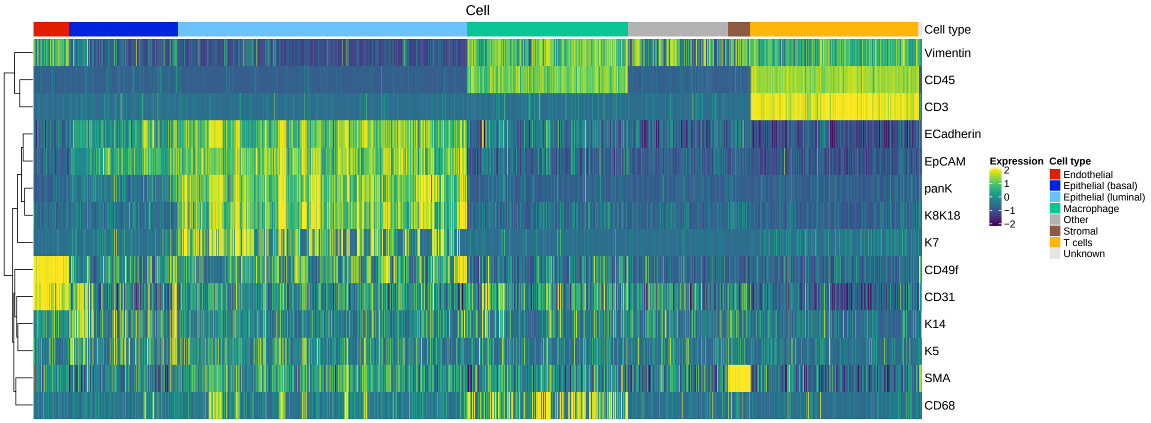


**Figure S7:** Expression heatmap for all marker proteins and the associated Astir cell types for Wagner.

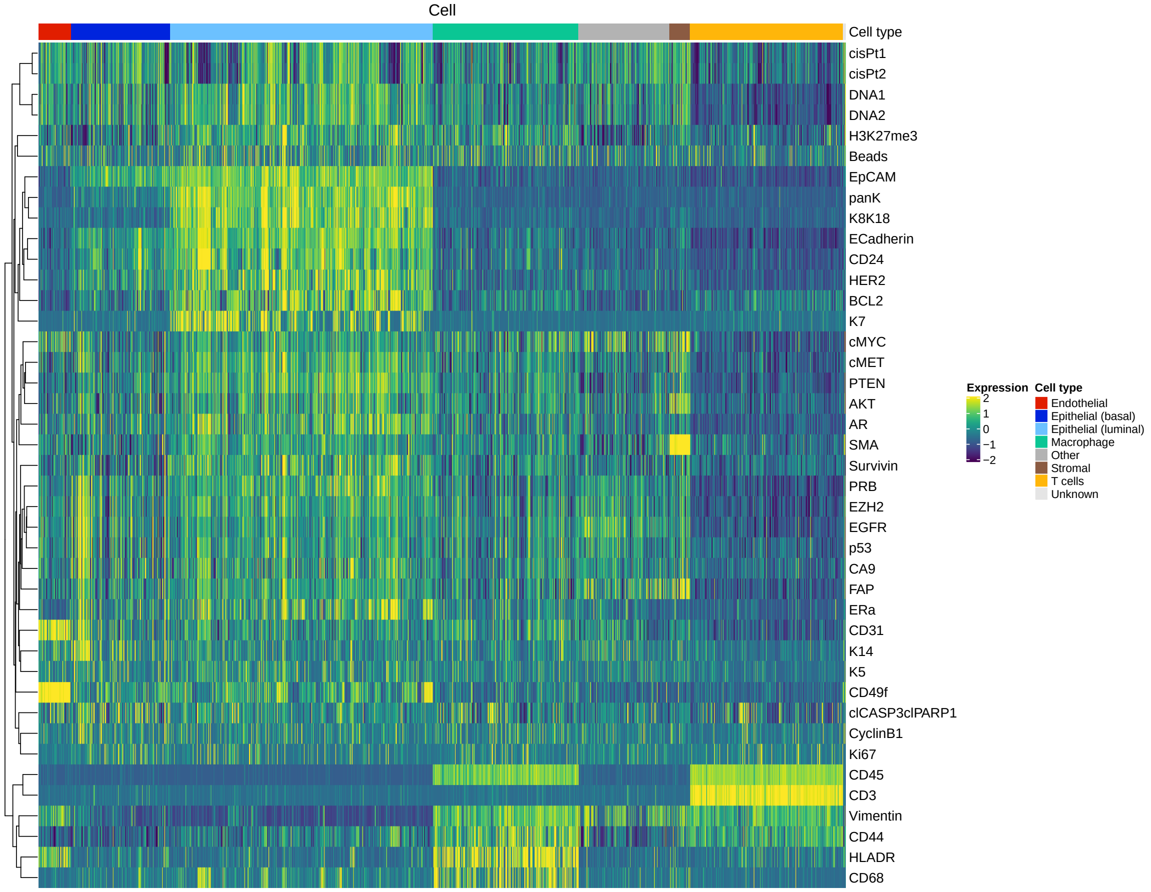


**Figure S8:** Expression heatmap for all proteins and the associated Astir cell types for Wagner.

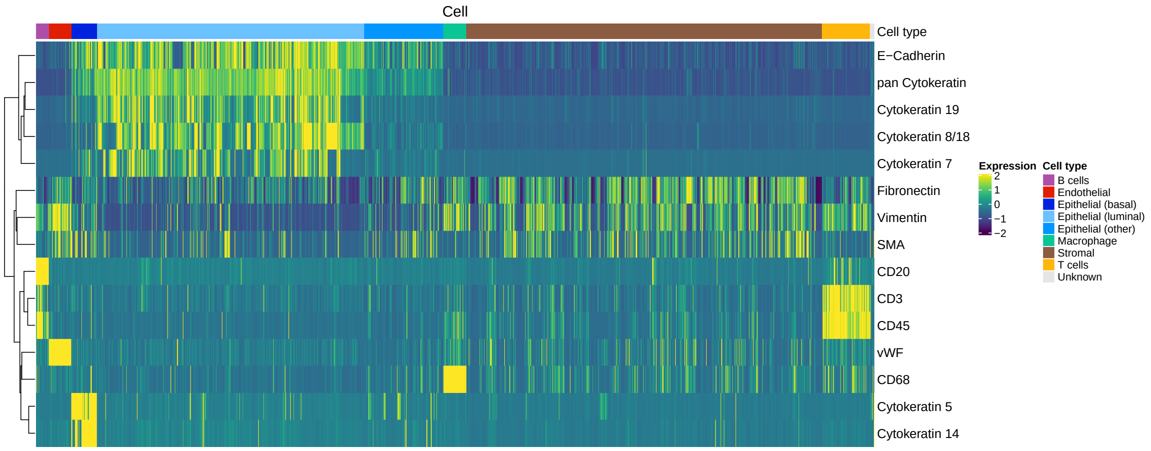


**Figure S9:** Expression heatmap for all marker proteins and the associated Astir cell types for Zurich.

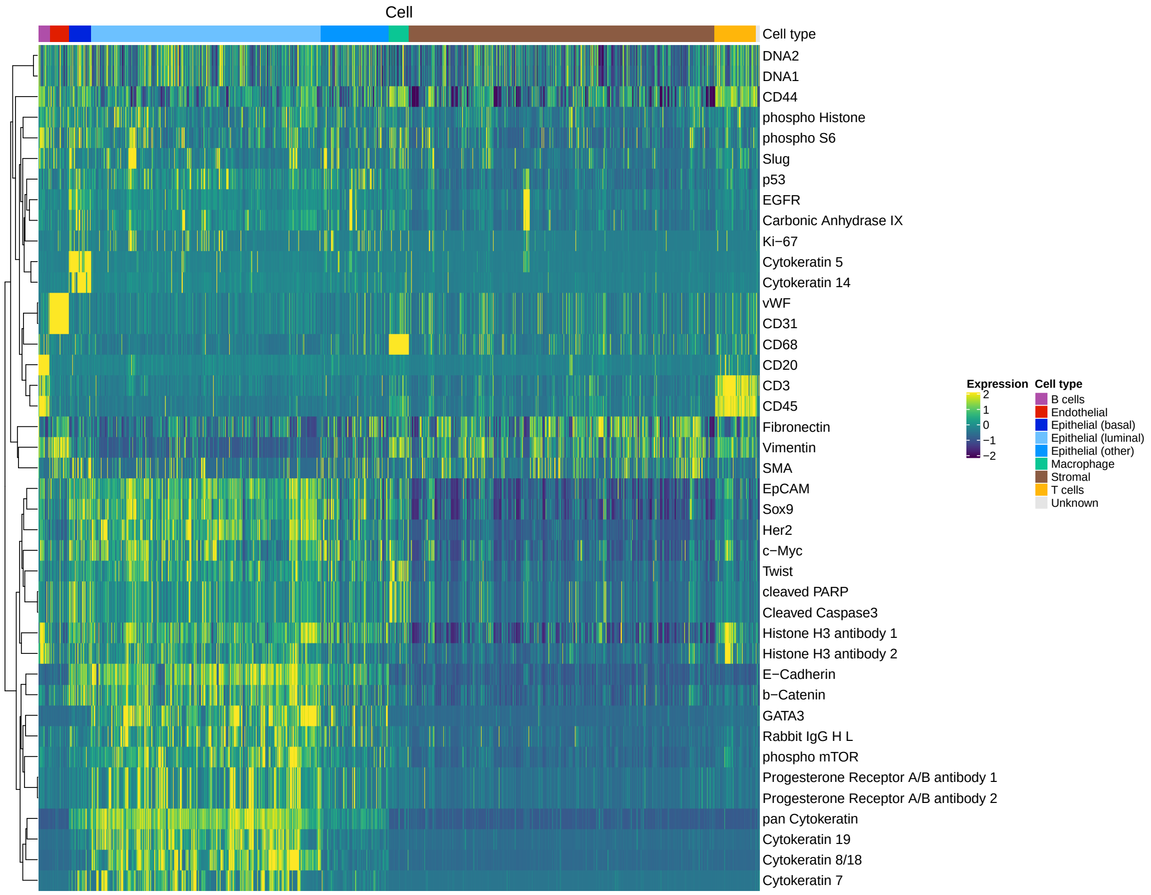


**Figure S10:** Expression heatmap for all proteins and the associated Astir cell types for Zurich.

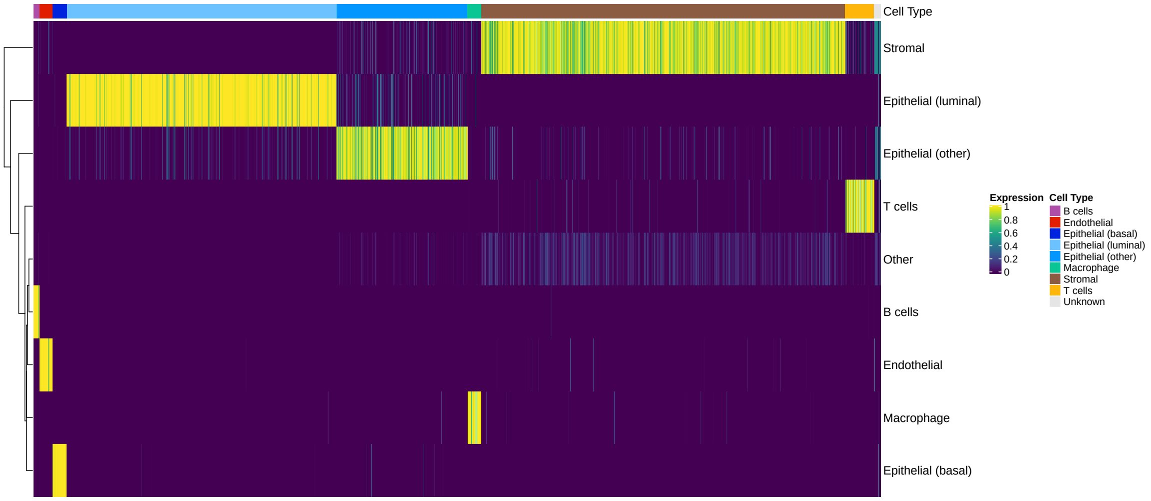


**Figure S11:** Heatmap depicting the cell type probabilities associated with each cell and Astir cell type for Basel.

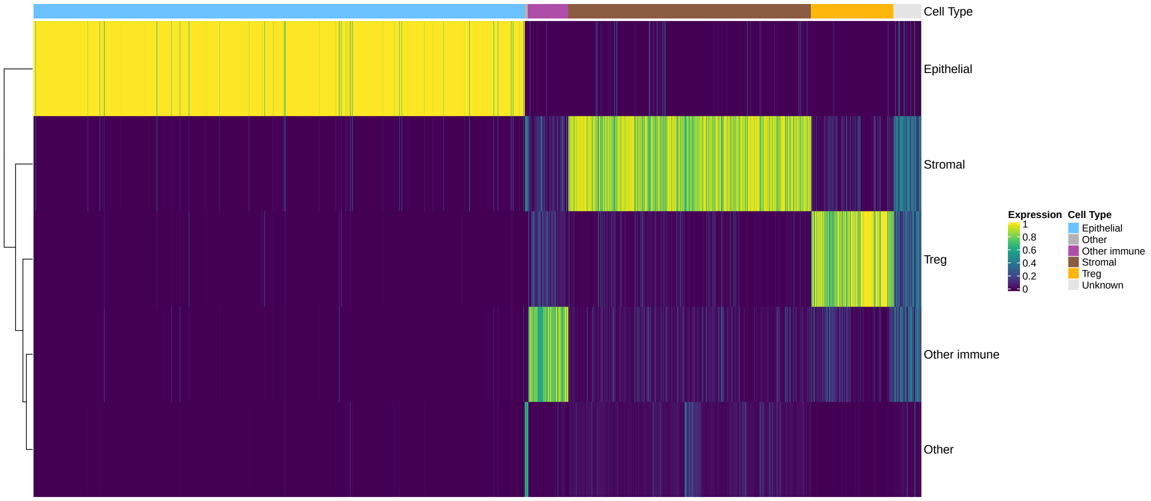


**Figure S12:** Heatmap depicting the cell type probabilities associated with each cell and Astir cell type for Lin.

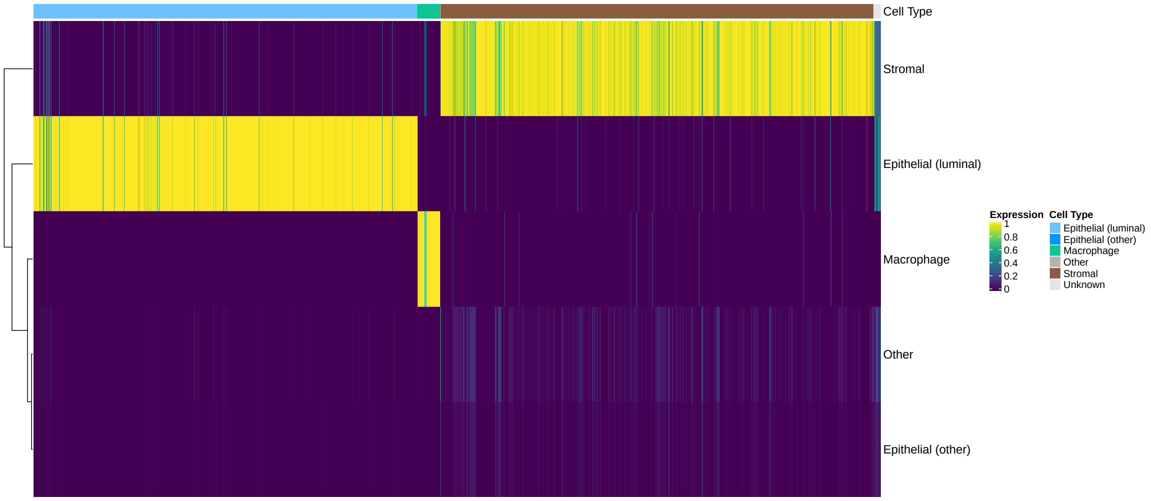


**Figure S13:** Heatmap depicting the cell type probabilities associated with each cell and Astir cell type for Schapiro.

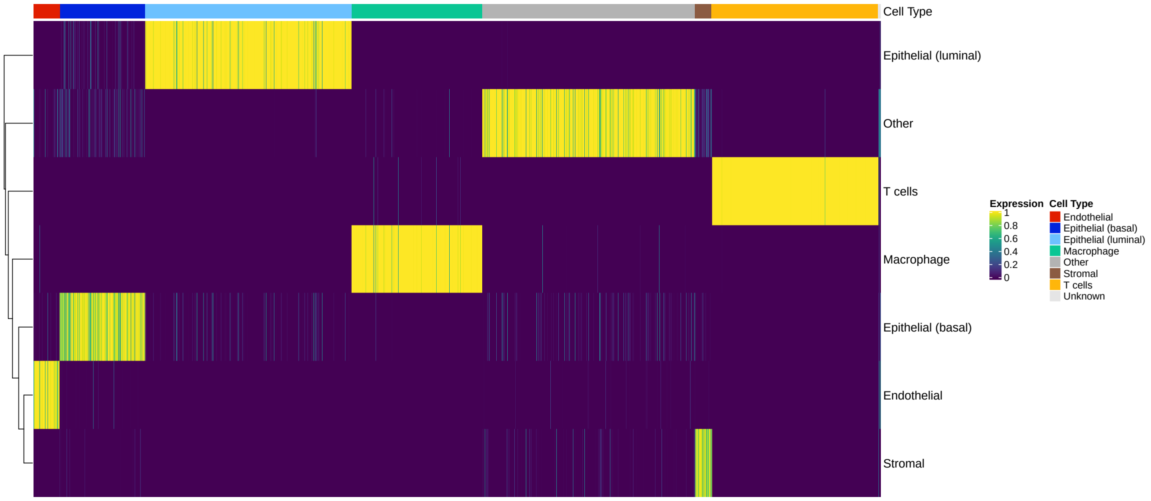


**Figure S14:** Heatmap depicting the cell type probabilities associated with each cell and Astir cell type for Wagner.

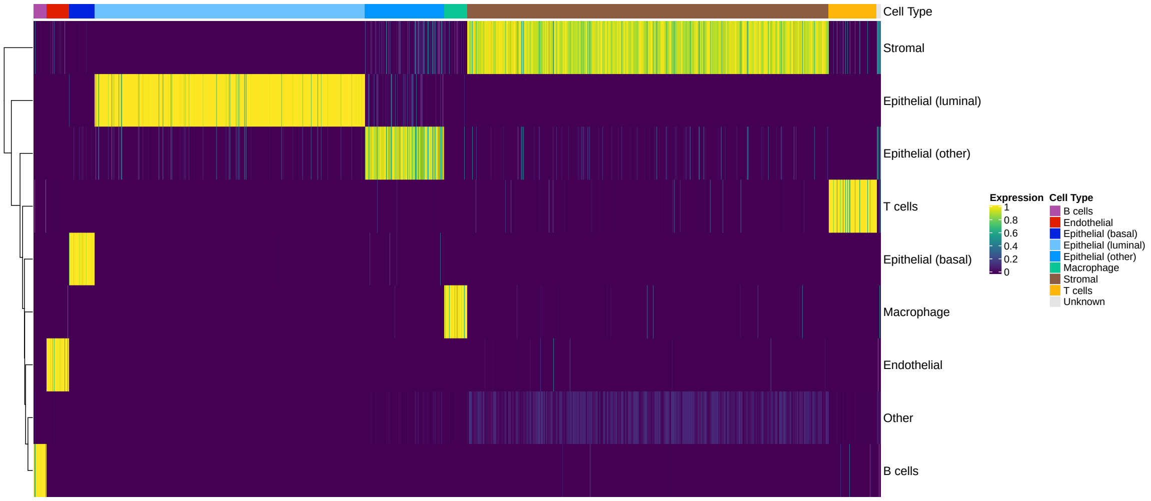


**Figure S15:** Heatmap depicting the cell type probabilities associated with each cell and Astir cell type for Zurich.

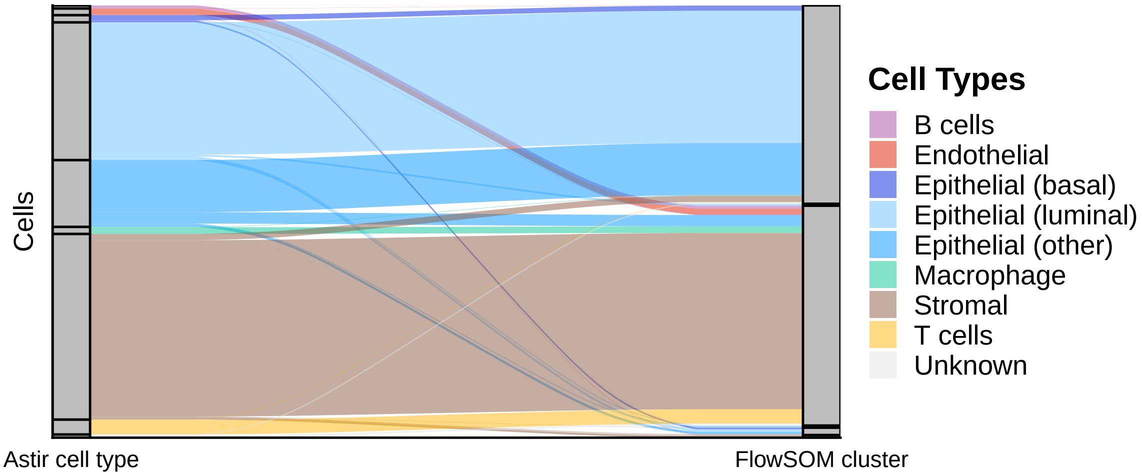


**Figure S16:** FlowSOM k7 specified markers Basel.

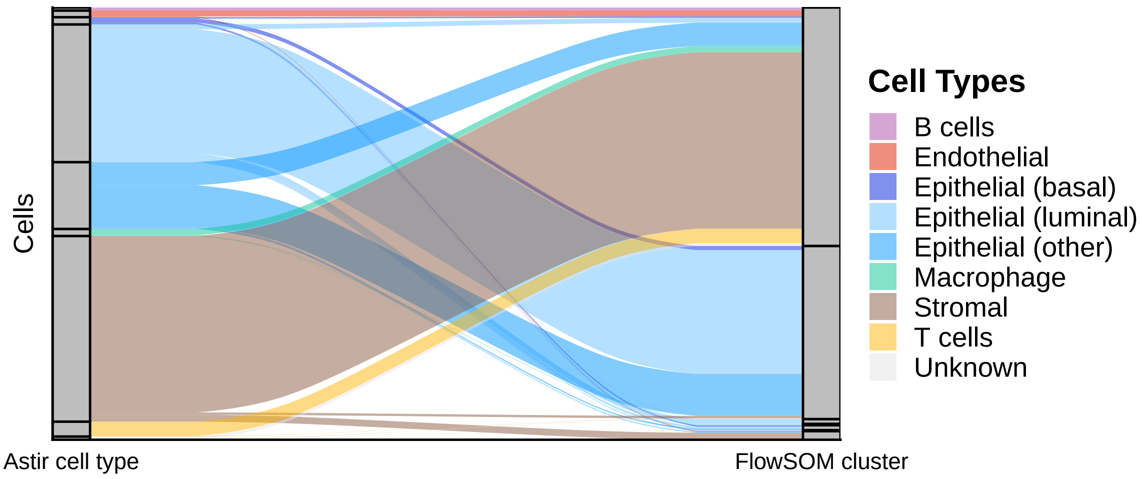


**Figure S17:** FlowSOM k7 all markers Basel.

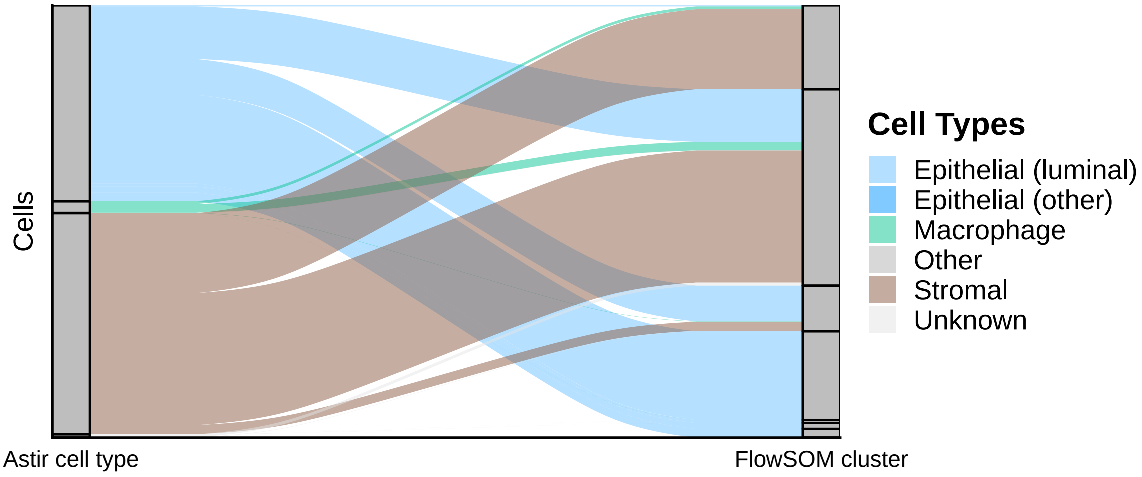


**Figure S18:** FlowSOM k7 specified markers Schapiro.

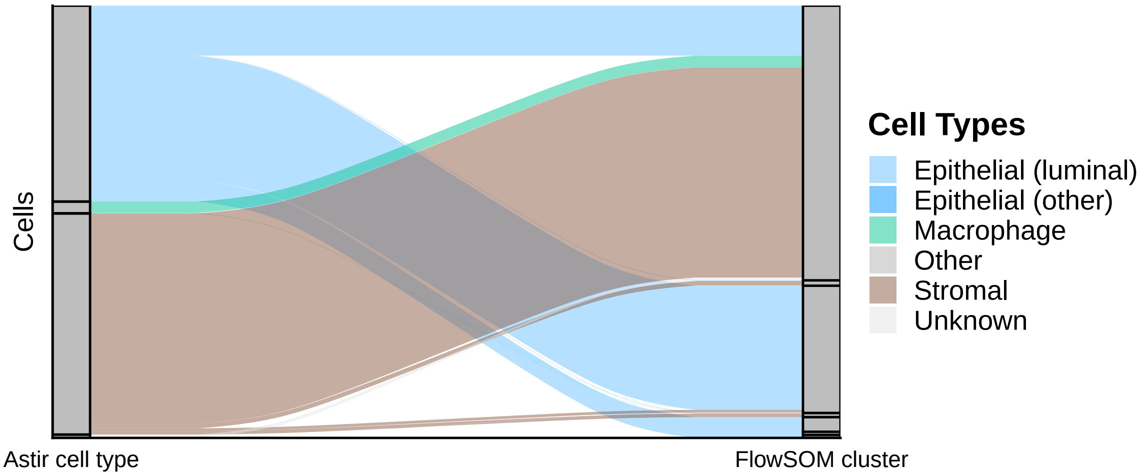


**Figure S19:** FlowSOM k7 specified markers Schapiro.

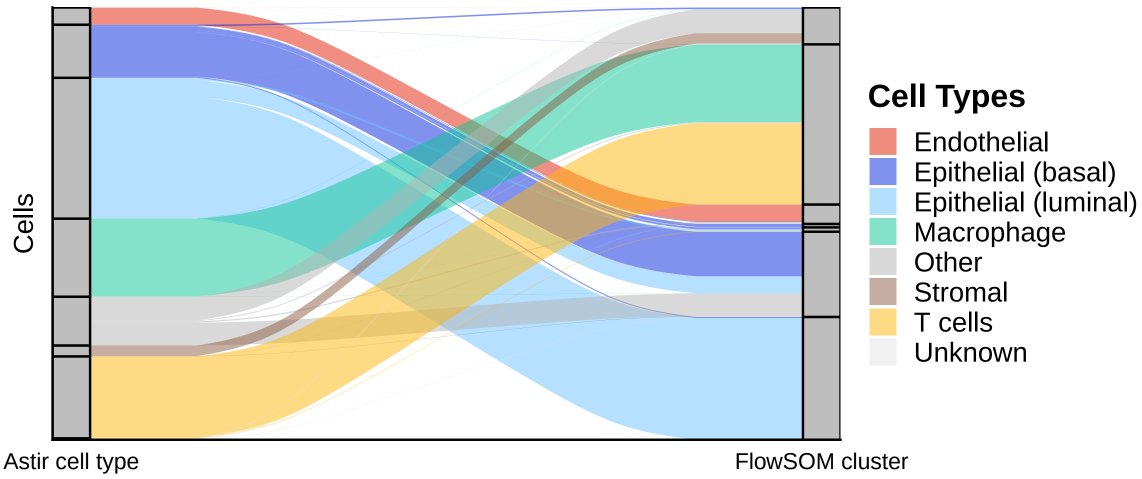


**Figure S20:** FlowSOM k7 specified markers Wagner.

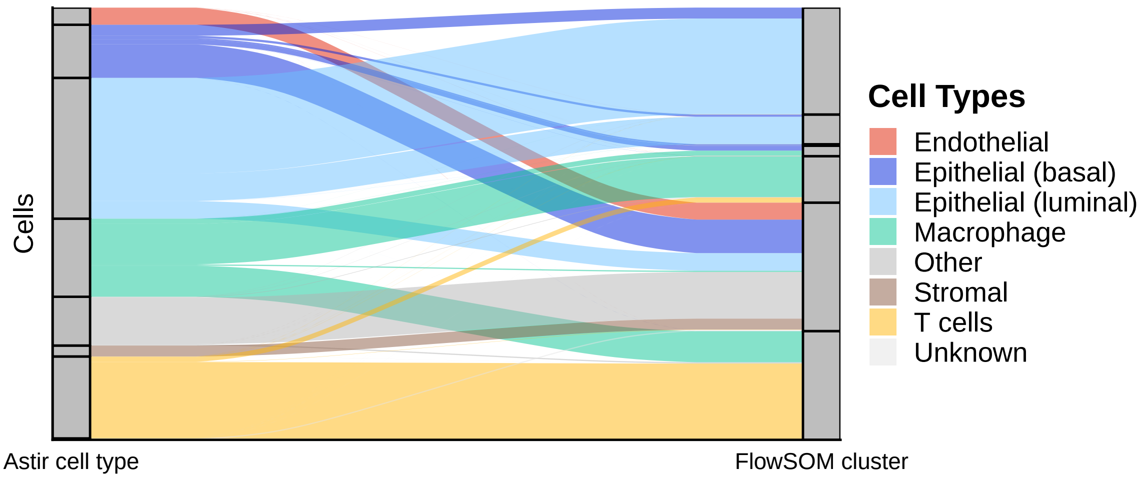


**Figure S21:** FlowSOM k7 all markers Wagner.

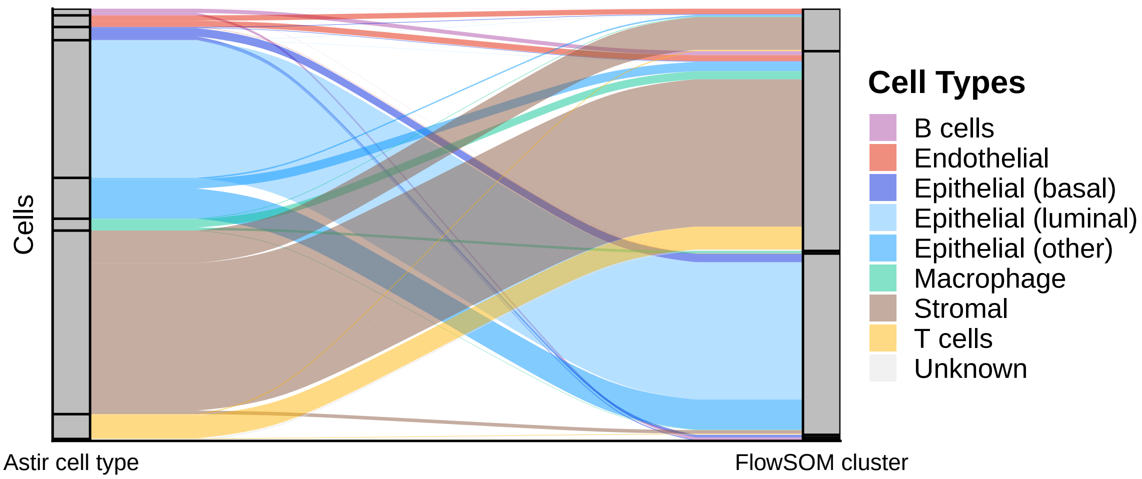


**Figure S22:** FlowSOM k7 specified markers Zurich.

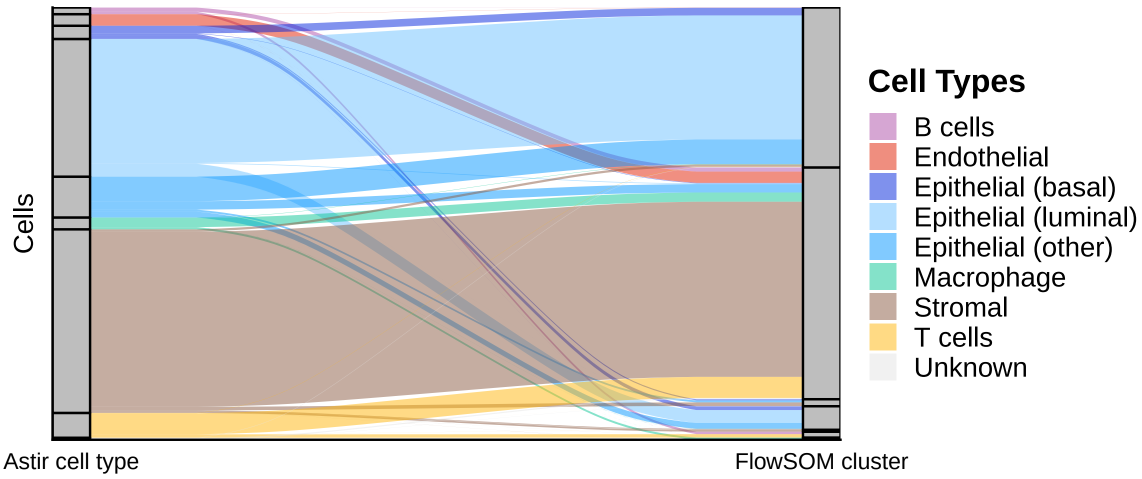


**Figure S23:** FlowSOM k7 all markers Zurich.

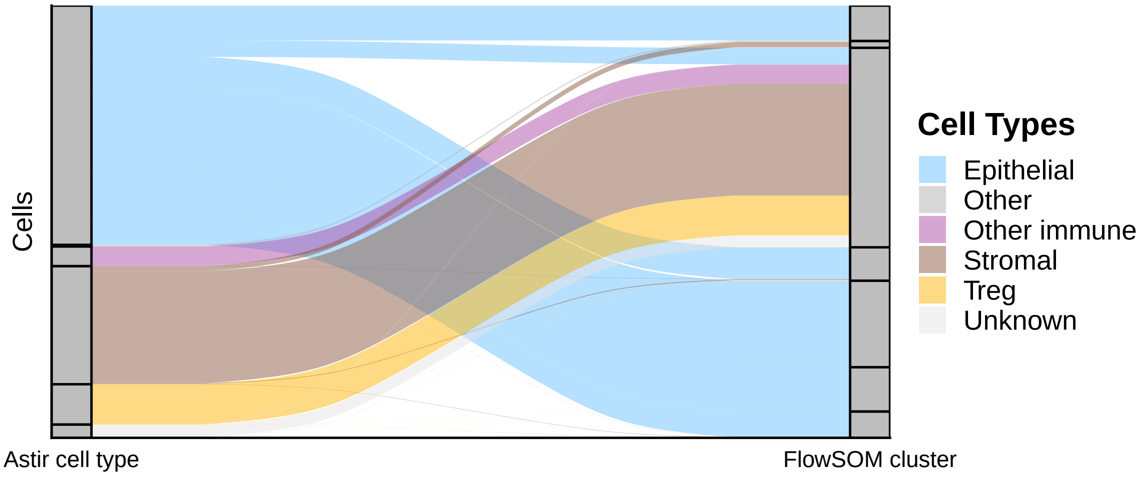


**Figure S24:** FlowSOM k7 specified markers Lin.

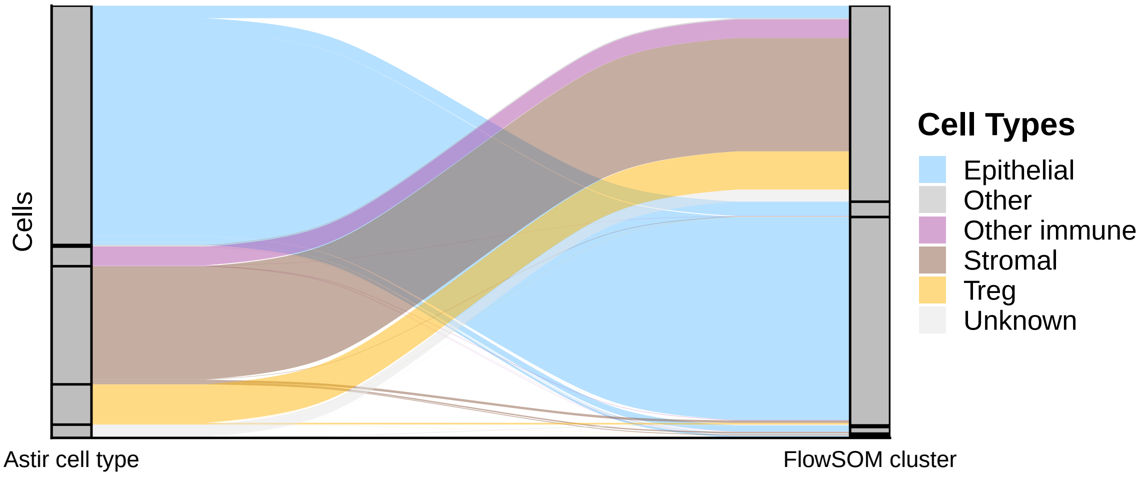


**Figure S25:** FlowSOM k7 all markers Lin.

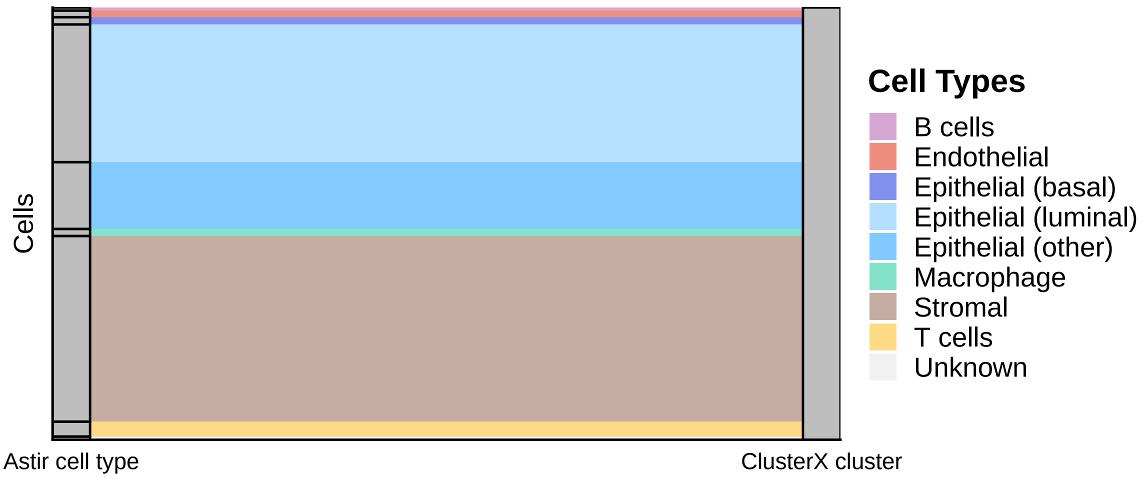


**Figure S26:** ClusterX specified markers Basel.

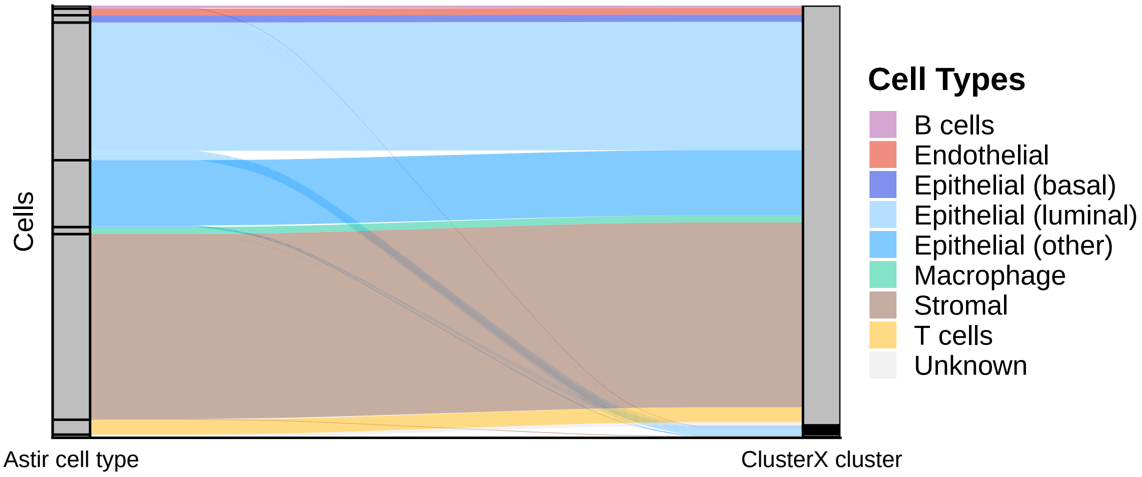


**Figure S27:** ClusterX all markers Basel.

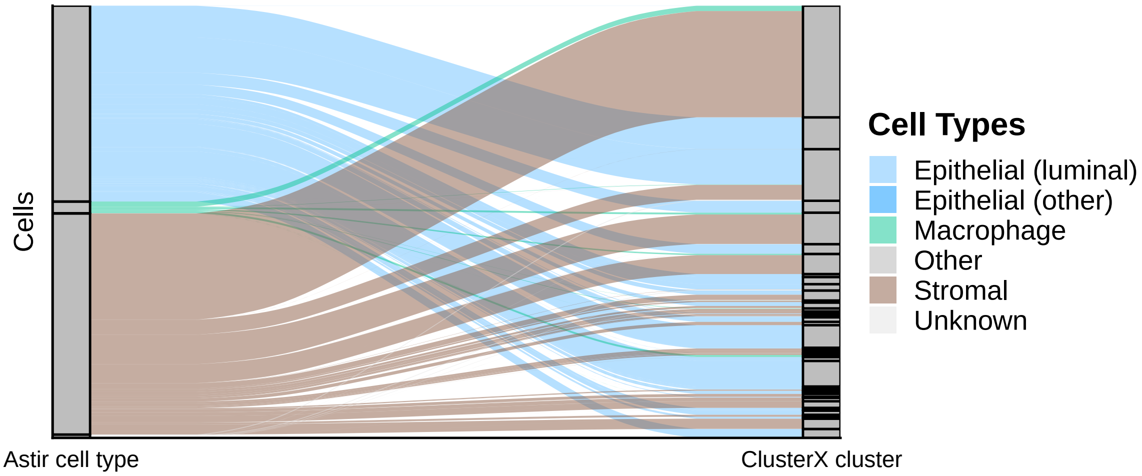


**Figure S28:** ClusterX specified markers Schapiro.

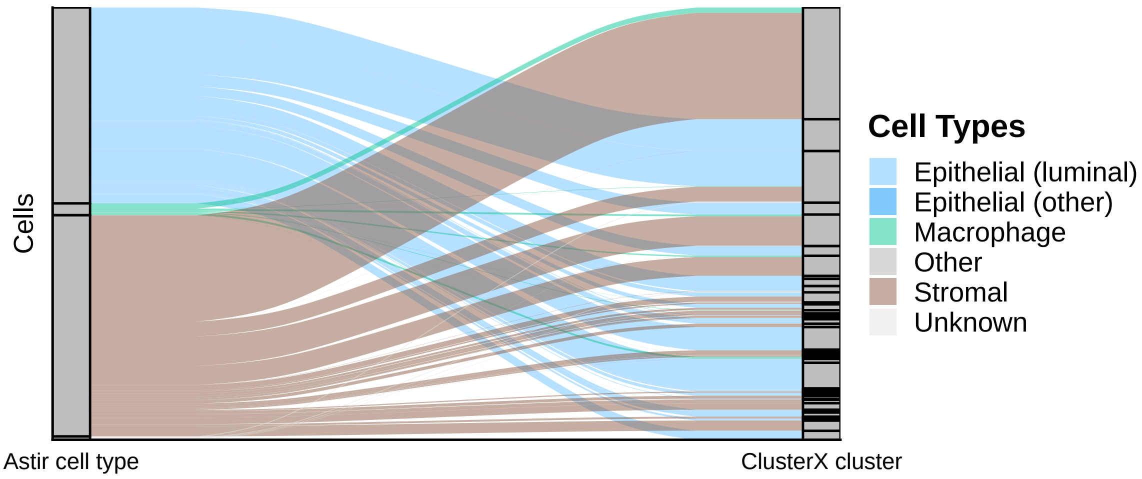


**Figure S29:** ClusterX all markers Schapiro.

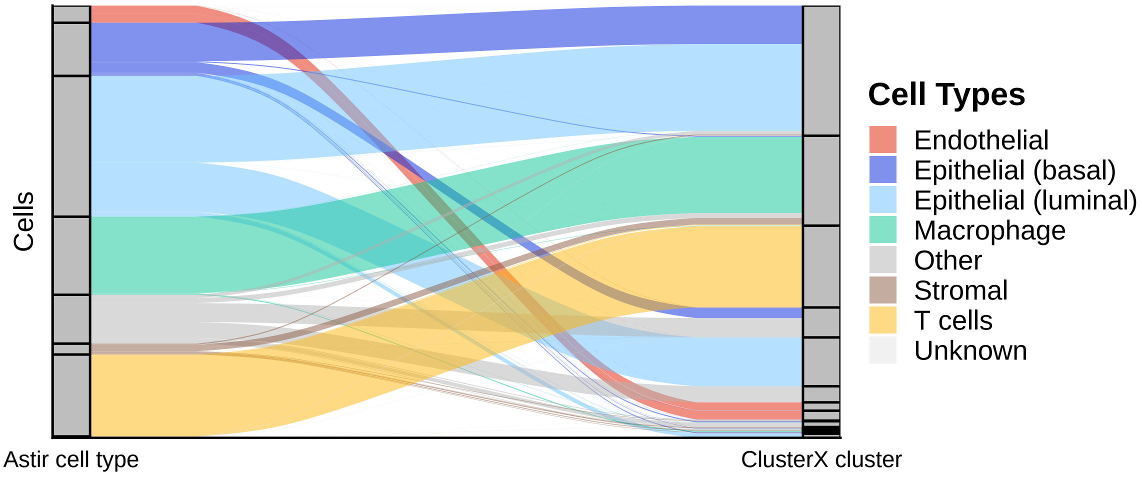


**Figure S30:** ClusterX specified markers Wagner.

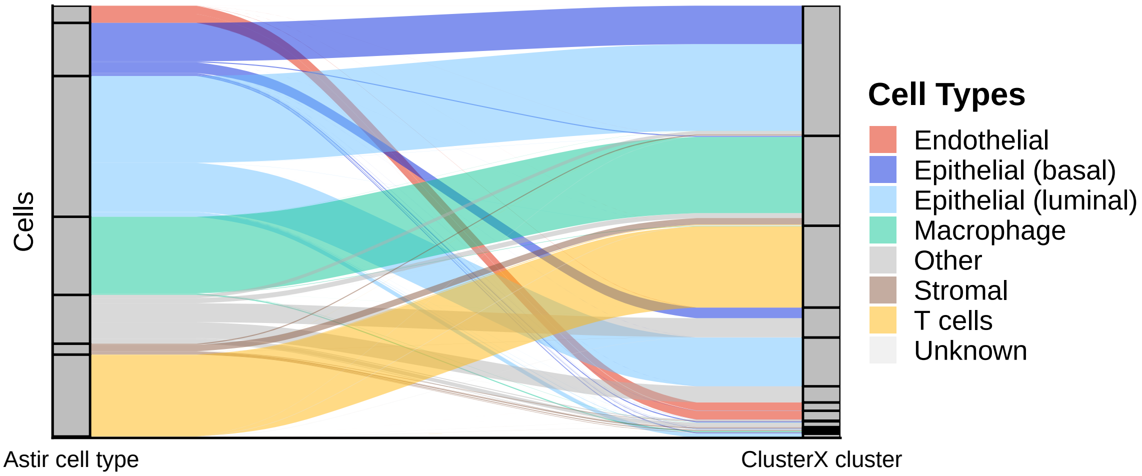


**Figure S31:** ClusterX all markers Wagner.

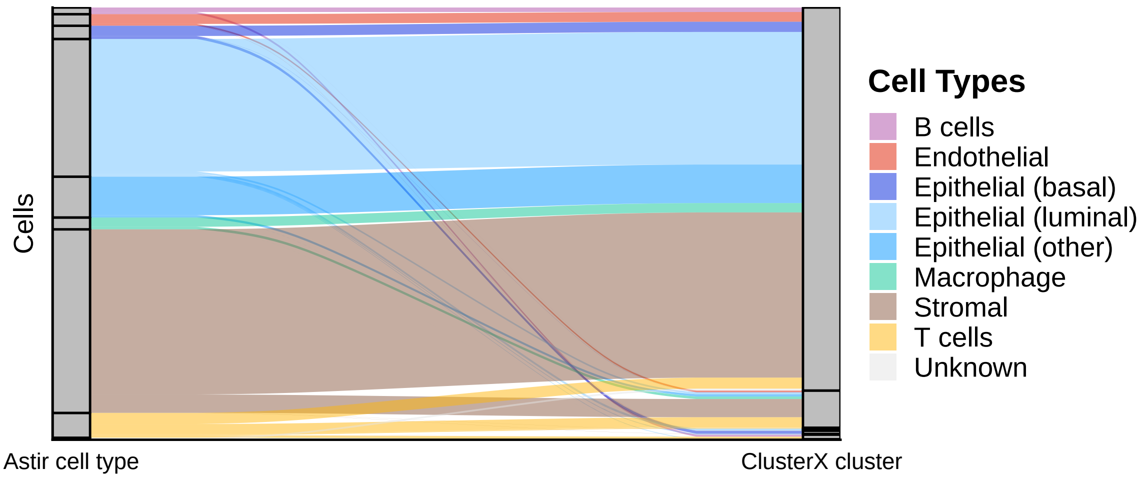


**Figure S32:** ClusterX specified markers Zurich.

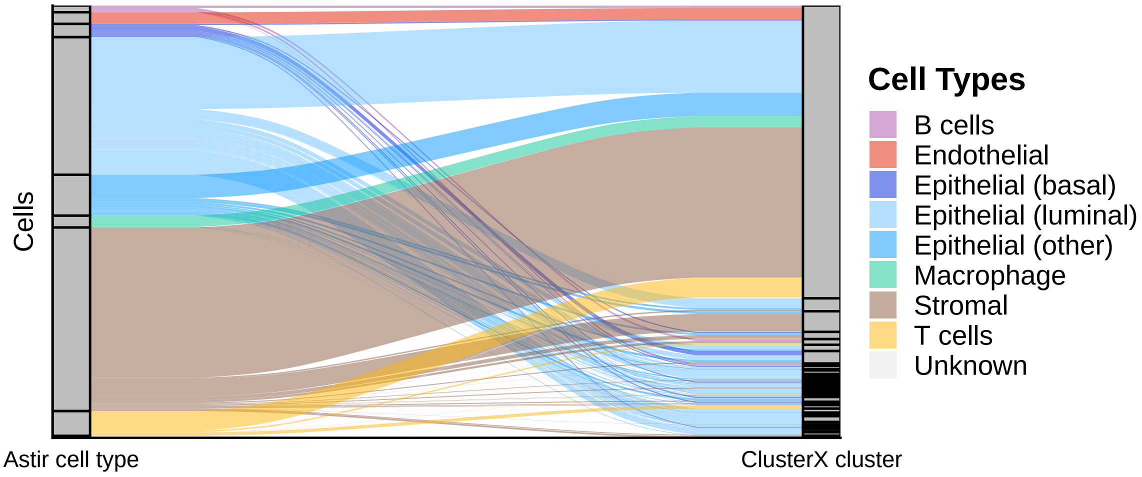


**Figure S33:** ClusterX all markers Zurich.

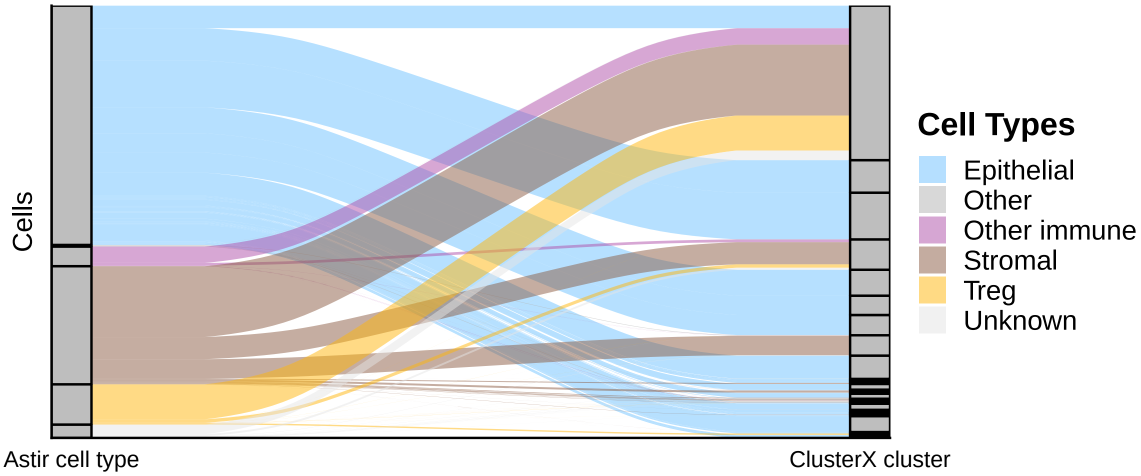


**Figure S34:** ClusterX specified markers Lin.

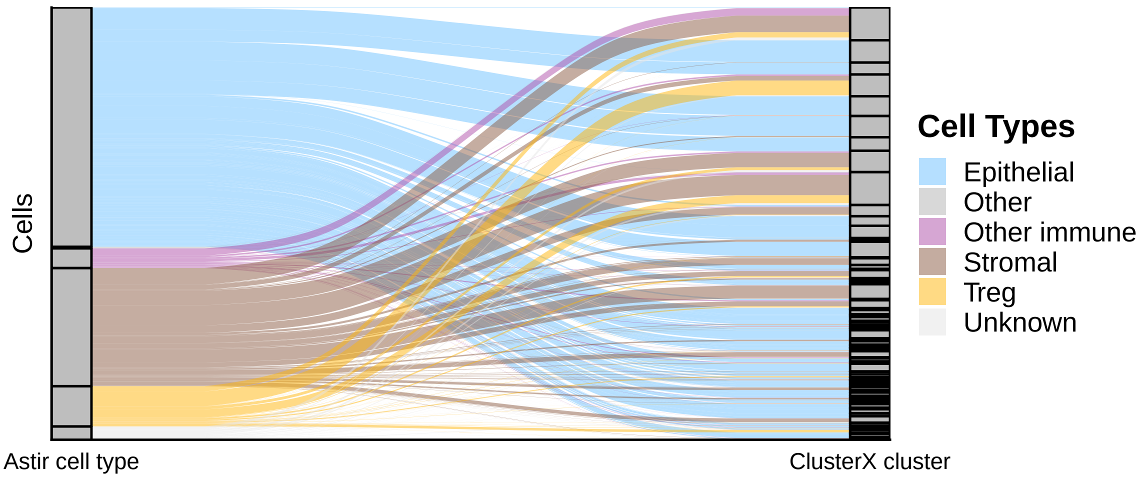


**Figure S35:** ClusterX all markers Lin.

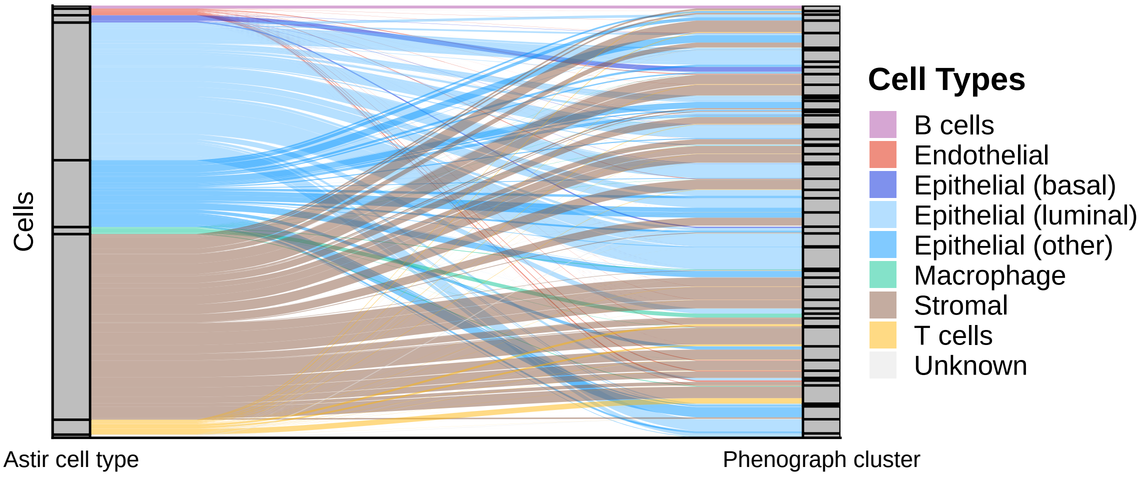


**Figure S36:** Phenograph specified markers Basel.

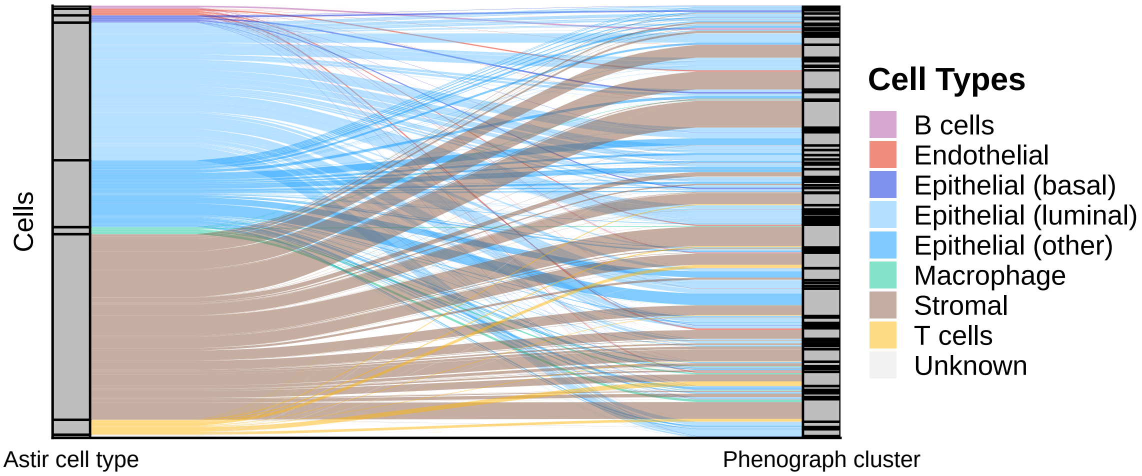


**Figure S37:** Phenograph all markers Basel.

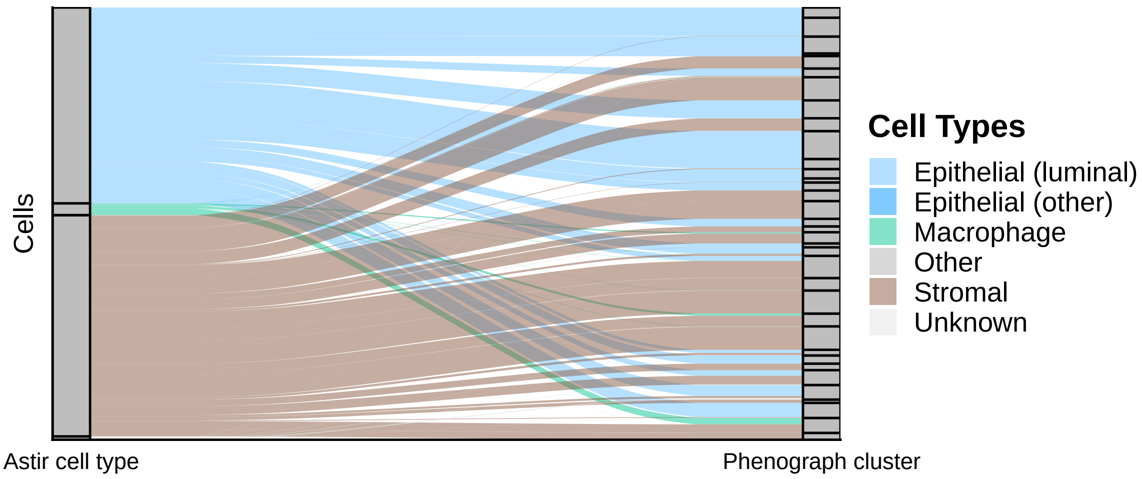


**Figure S38:** Phenograph specified markers Schapiro.

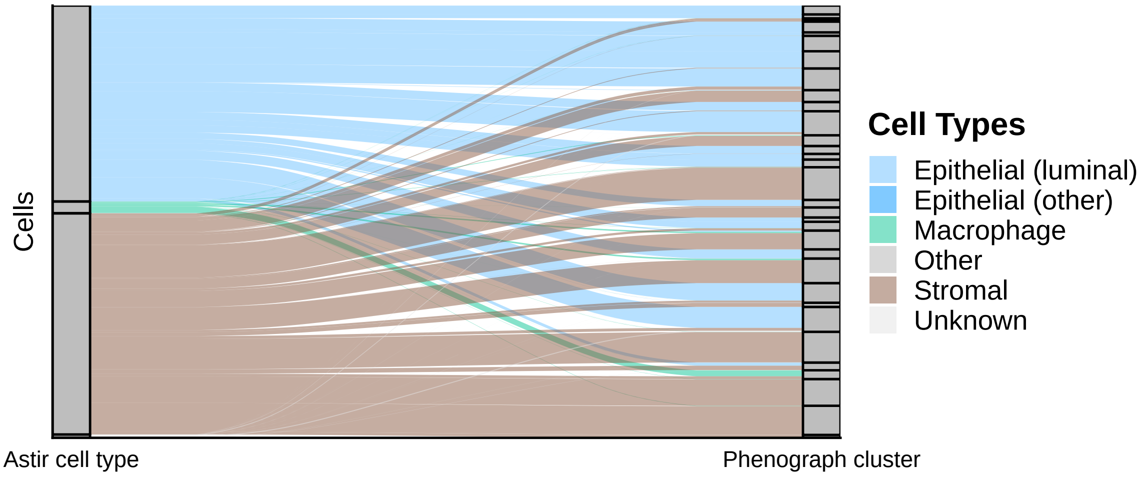


**Figure S39:** Phenograph all markers Schapiro.

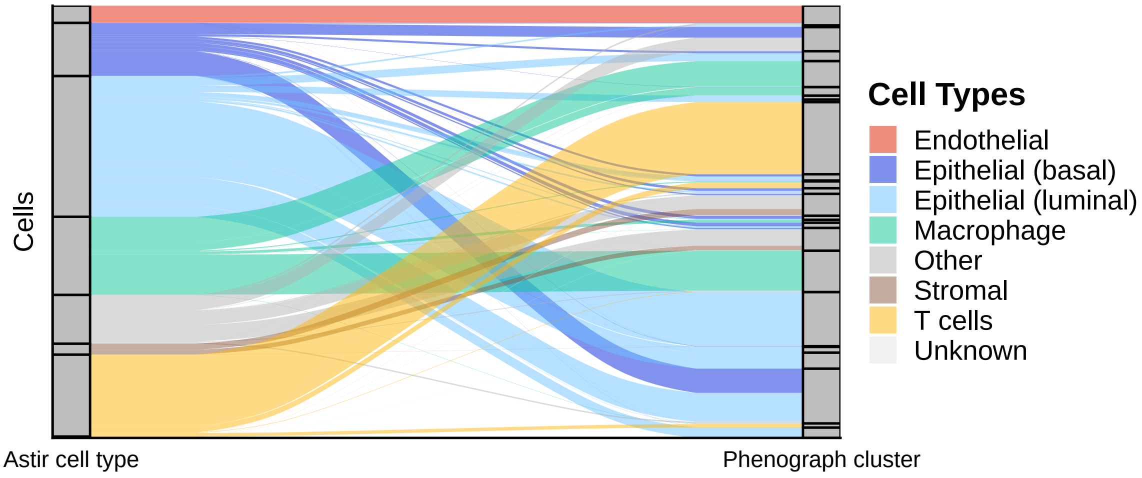


**Figure S40:** Phenograph specified markers Wagner.

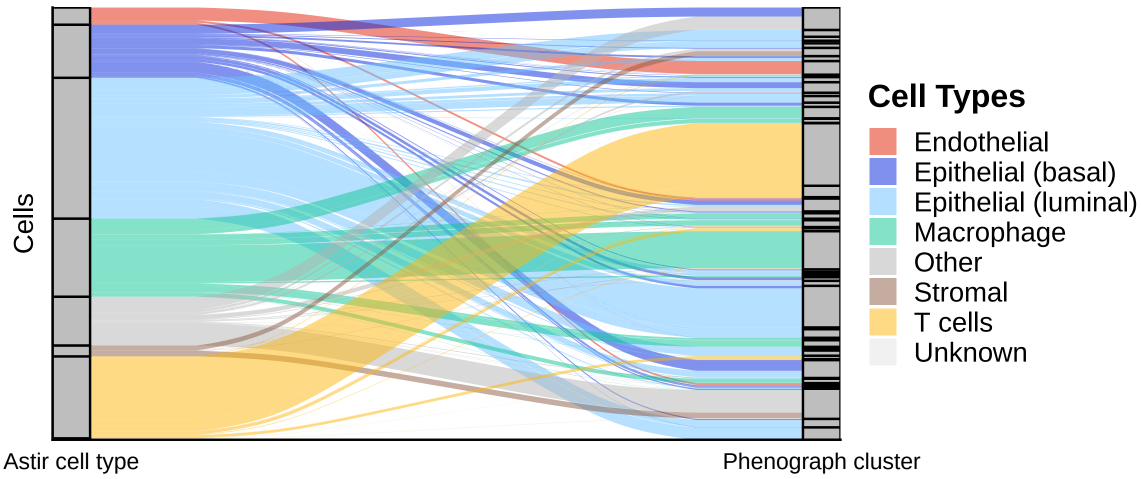


**Figure S41:** Phenograph all markers Wagner.

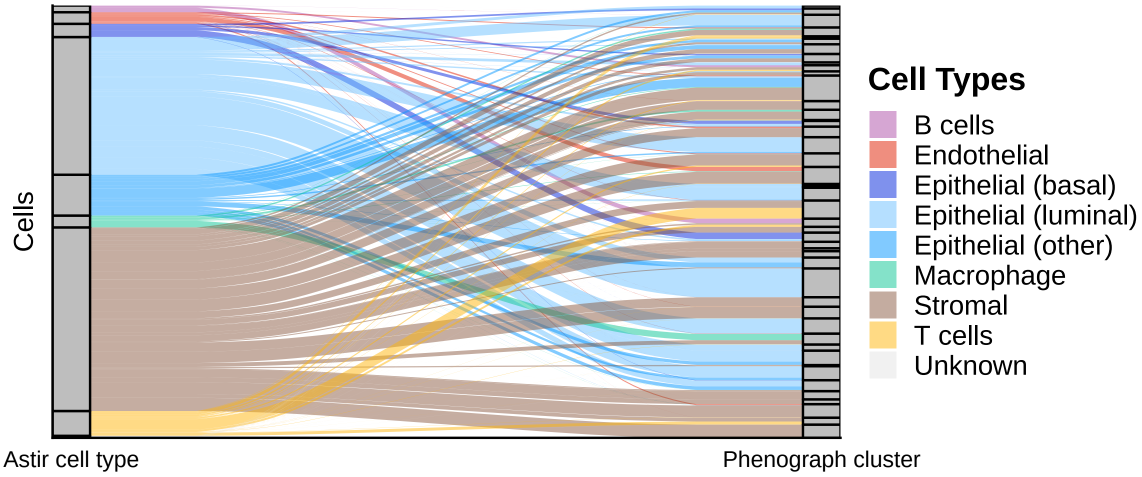


**Figure S42:** Phenograph specified markers Zurich.

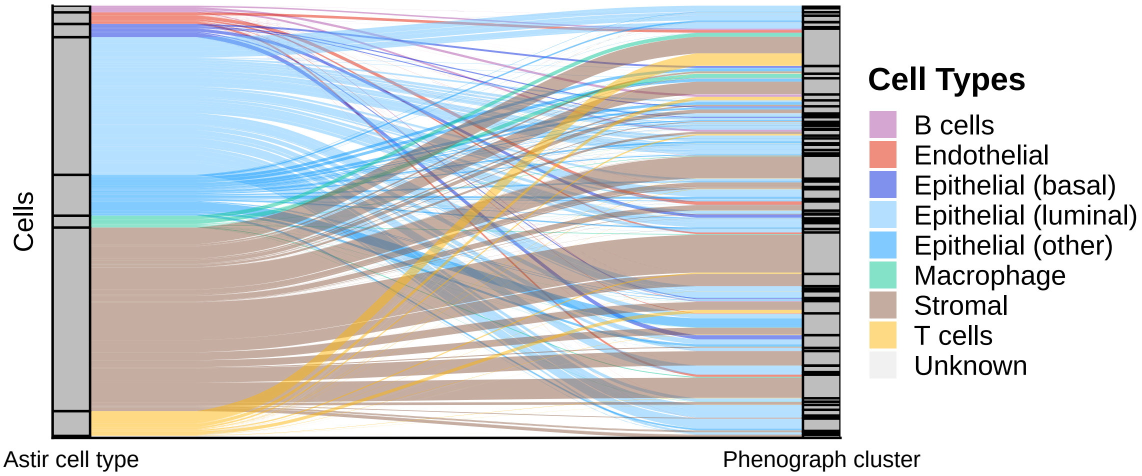


**Figure S43:** Phenograph all markers Zurich.

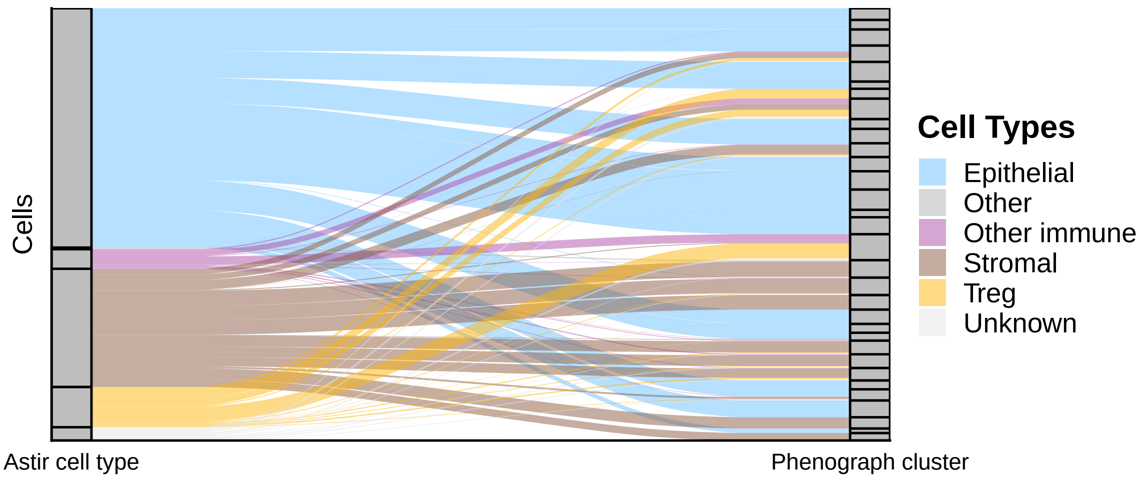


**Figure S44:** Phenograph specified markers Lin.

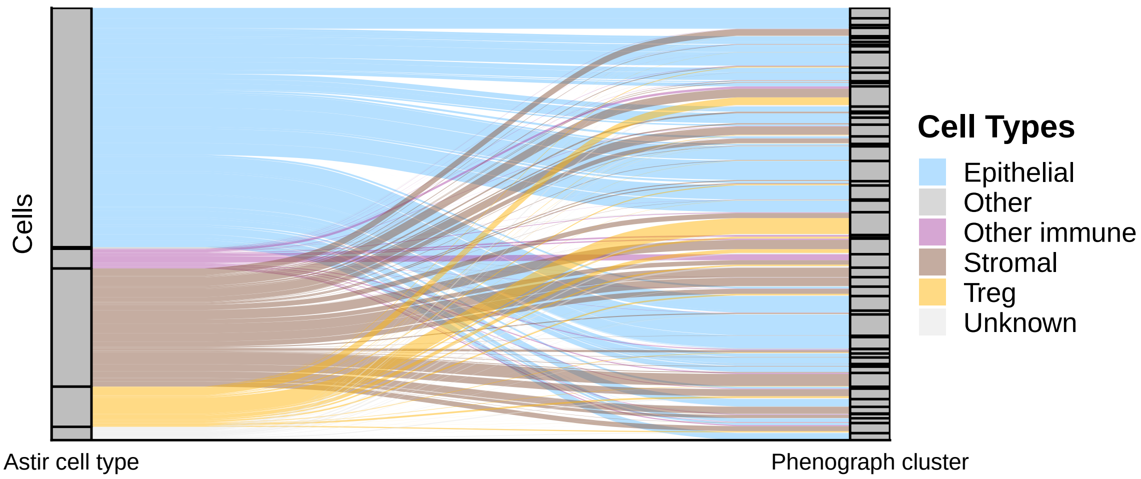


**Figure S45:** Phenograph all markers Lin.

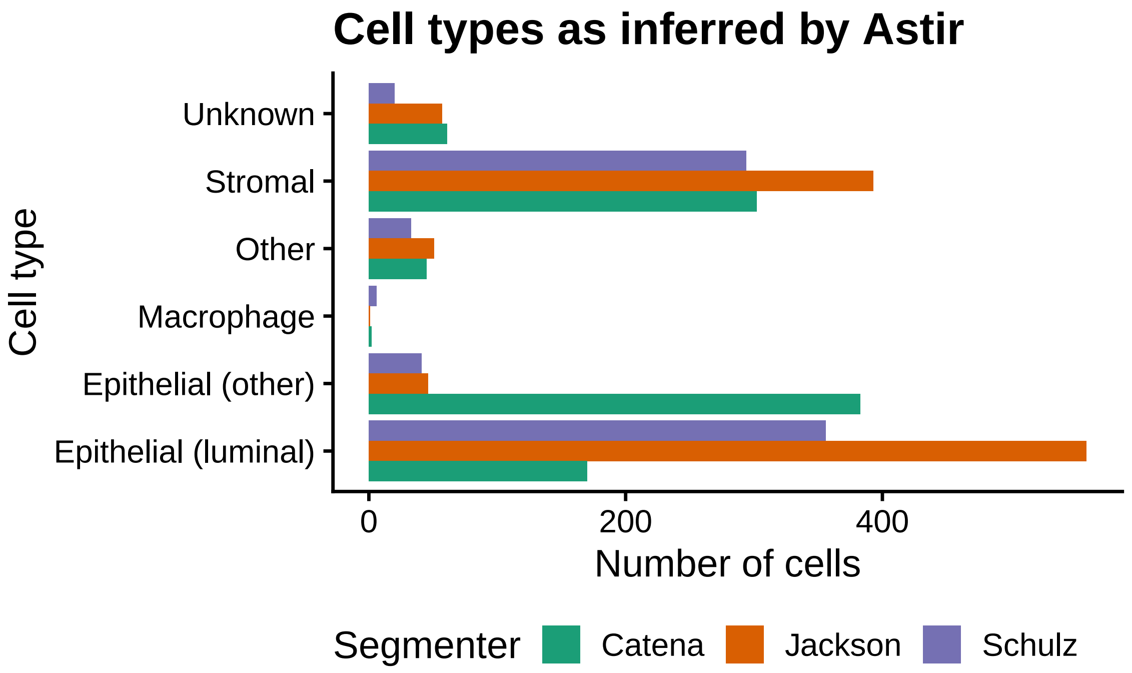


**Figure S46:** Number of cells inferred for each type and segmenter for the `Cy1x6_33` ROI.

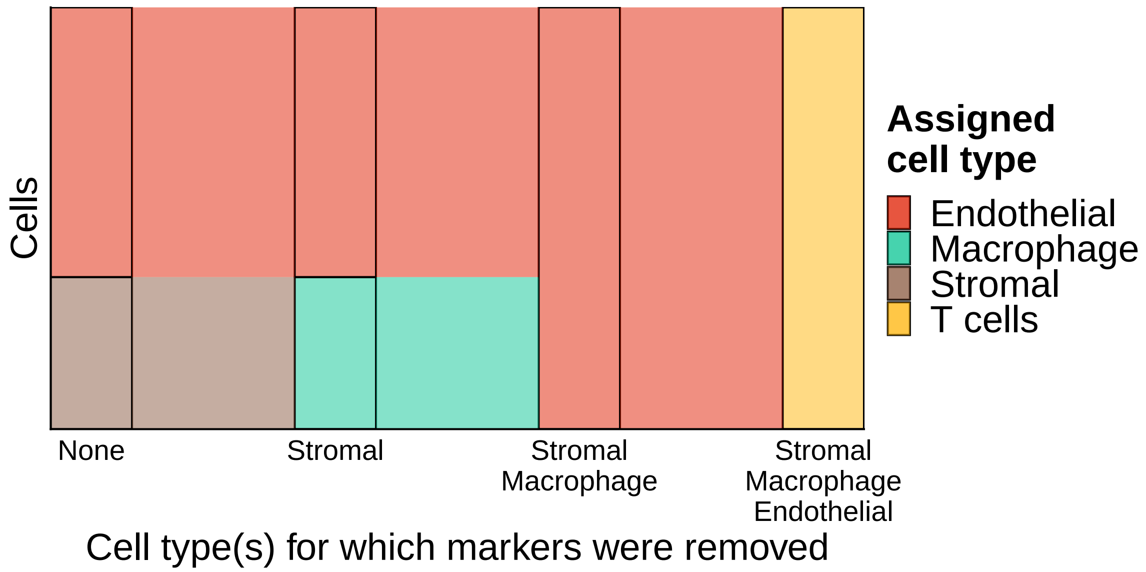


**Figure S47:** Cell type assignments for FlowSOM k8 with all markers in the Basel cohort upon removal of marker genes.

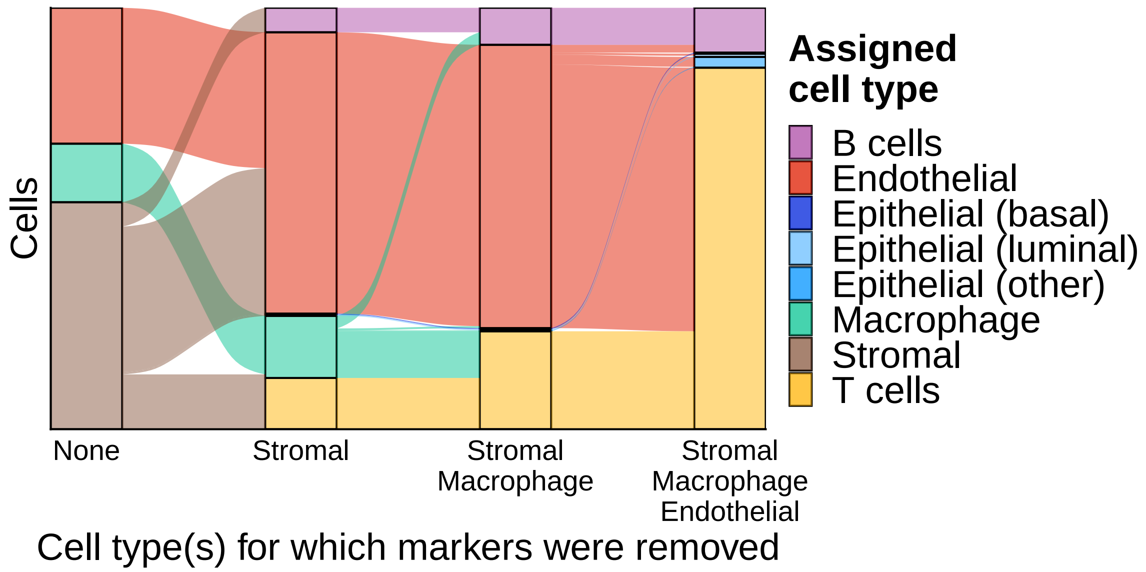


**Figure S48:** Cell type assignments for Phenograph k20 with all markers in the Basel cohort upon removal of marker genes.

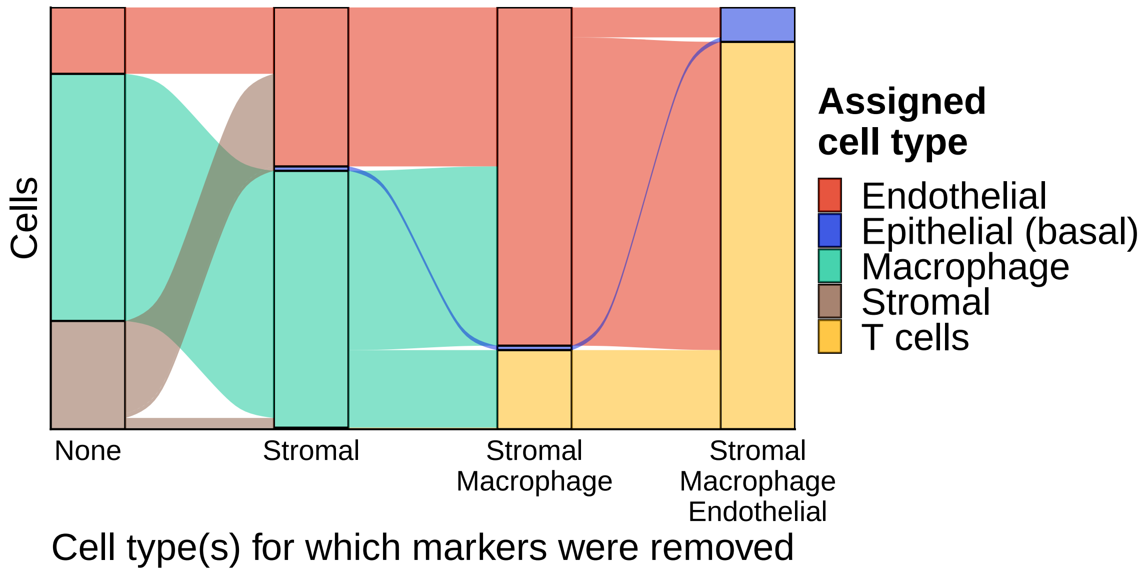


**Figure S49:** Cell type assignments for ClusterX with all markers in the Wagner cohort upon removal of marker genes.

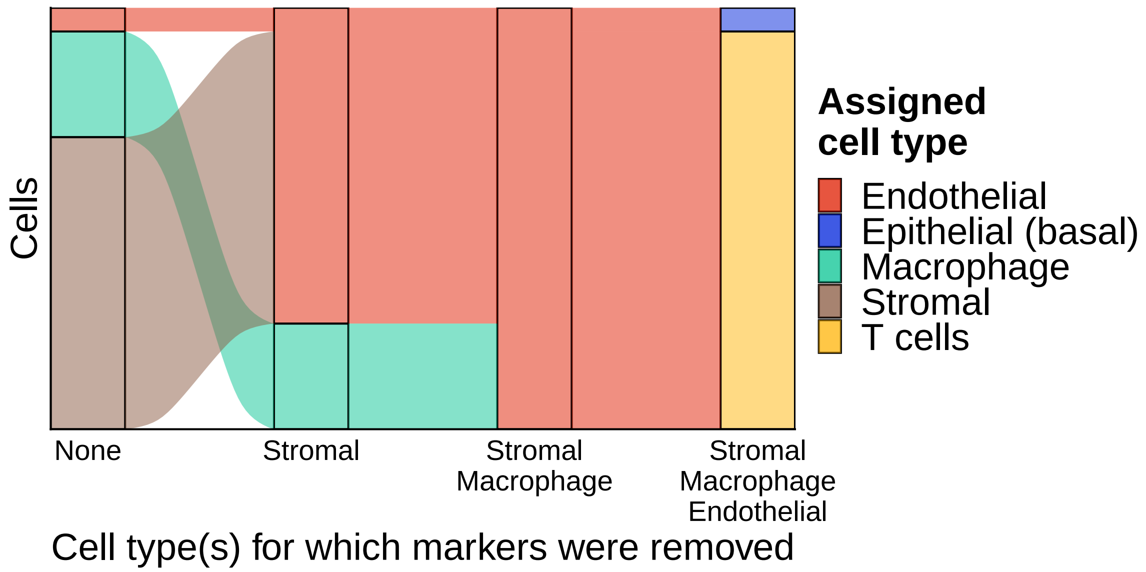


**Figure S50:** Cell type assignments for FlowSOM k7 with all markers in the Wagner cohort upon removal of marker genes.

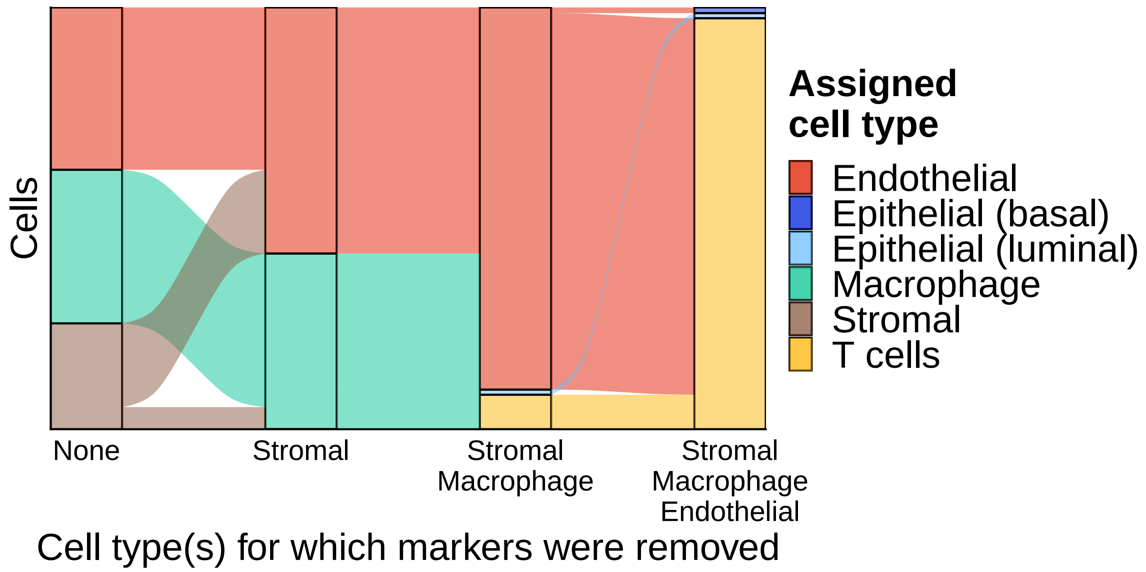


**Figure S51:** Cell type assignments for Phenograph k20 with all markers in the Wagner cohort upon removal of marker genes.

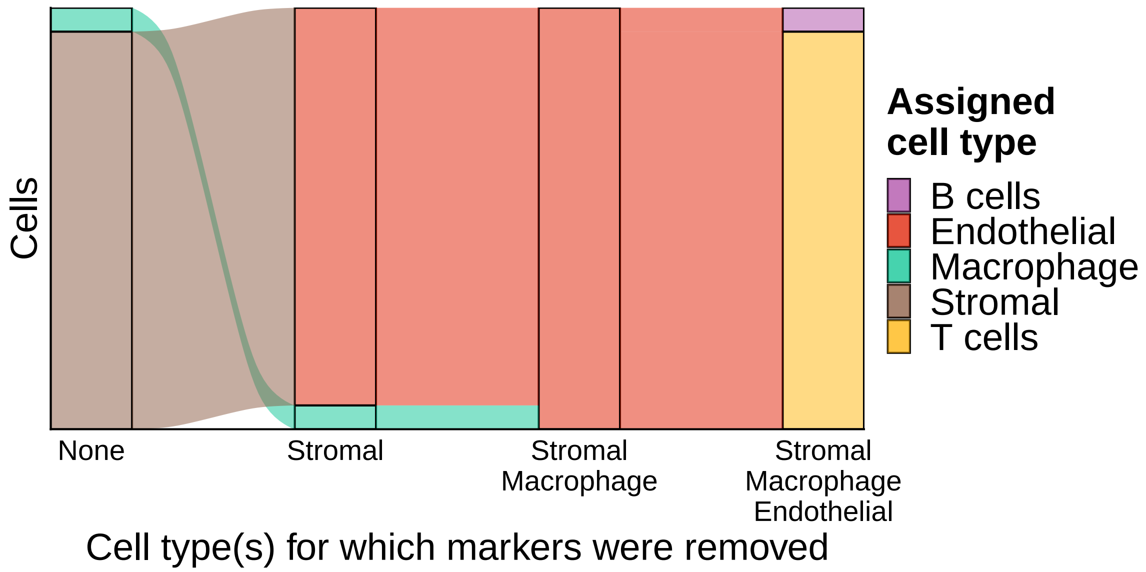


**Figure S52:** Cell type assignments for FlowSOM k7 with specified markers in the Zurich cohort upon removal of marker genes.

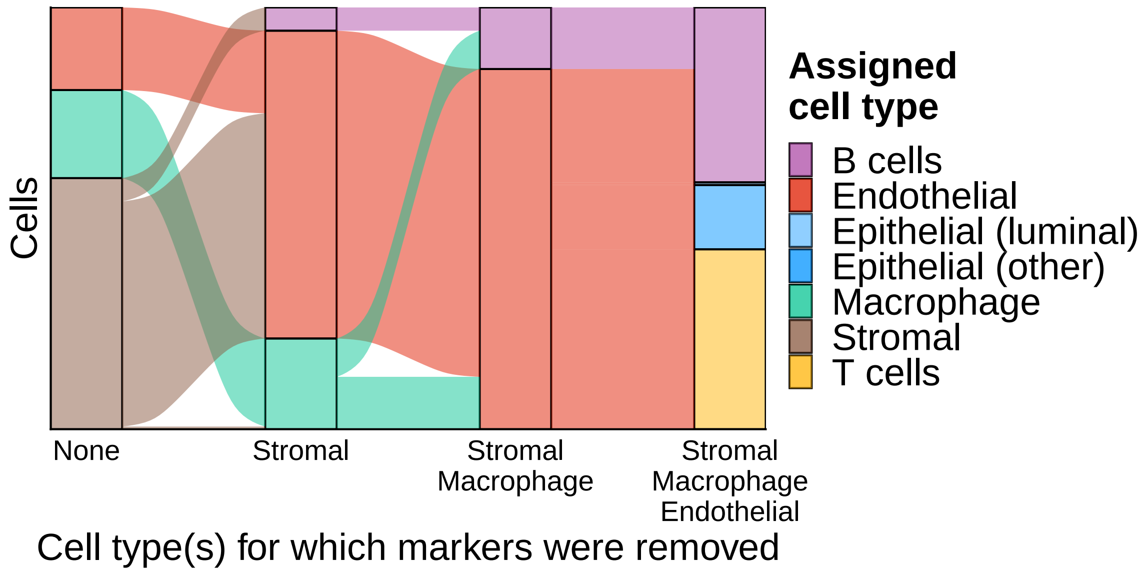


**Figure S53:** Cell type assignments for Phenograph k20 with specified markers in the Zurich cohort upon removal of marker genes.

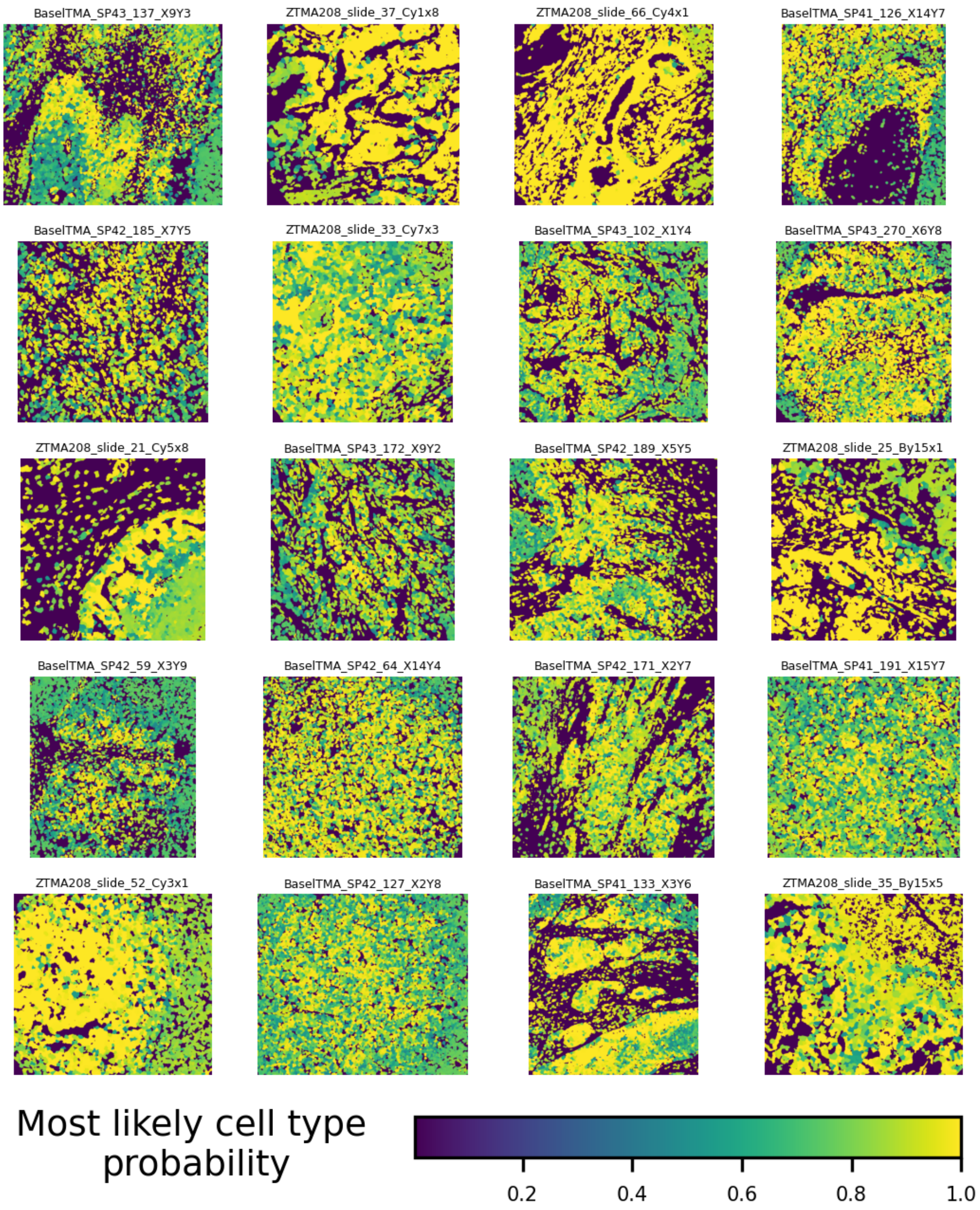


**Figure S54:** Representative cores from the Zurich and Basel cohorts showing the maximum probability associated with each cell.
